# Local and population-level responses of Greater sage-grouse to oil and gas development and climatic variation in Wyoming

**DOI:** 10.1101/028274

**Authors:** Rob R. Ramey, Joseph L. Thorley, Alexander S. Ivey

## Abstract

**Background:** Spatial scale is important when studying ecological processes. The Greater sage-grouse (*Centrocercus urophasianus*) is a large sexually dimorphic tetraonid that is endemic to the sagebrush biome of western North America. The impacts of oil and gas development at individual leks has been well-documented. However, no previous studies have quantified the population-level response.

**Methods:** Hierarchical models were used to estimate the effects of the areal disturbance due to well pads as well as climatic variation on individual lek counts and Greater sage-grouse populations (management units) over 32 years. The lek counts were analyzed using General Linear Mixed Models while the management units were analyzed using Gompertz Population Dynamic Models. The models were fitted using frequentist and Bayesian methods. An information-theoretic approach was used to identify the most important spatial scale and time lags. The relative importance of oil and gas and climate at the local and population-level scales was assessed using information-theoretic (Akaike’s weights) and estimation (effect size) statistics.

**Results:** At the local scale, oil and gas was an important negative predictor of the lek count. At the population scale, there was only weak support for oil and gas as a predictor of density changes but the estimated impacts on the long-term carrying capacity were consistent with summation of the local impacts. Regional climatic variation, as indexed by the Pacific Decadal Oscillation, was an important positive predictor of density changes at both the local and population-level (particularly in the most recent part of the time series).

**Conclusions:** Additional studies to reduce the uncertainty in the range of possible effects of oil and gas at the population scale are required. Wildlife agencies need to account for the effects of regional climatic variation when managing sage-grouse populations.

## INTRODUCTION

> If we study a system at an inappropriate scale, we may not detect its actual dynamics and patterns but may instead identify patterns that are artifacts of scale. Because we are clever at devising explanations of what we see, we may think we understand the system when we have not even observed it correctly.
>
> — Wiens (1989)

Effective conservation of a species requires an understanding of how human activities influence its distribution and abundance. Although much of science proceeds by experimental studies to understand the causal links between actions and responses, ethical, practical and statistical considerations typically prevent population-level experiments on species of concern. Consequently, many conservation-based ecological studies are forced to infer the population-level consequences of anthropogenic alterations from local gradients (Fukami and Wardle, 2005) in density (Gill et al., 2001), movement, habitat use, physiology, genetics, reproductive success or survival. However, local gradients may not accurately predict the population-level response (Fodrie et al., 2014).

The Greater sage-grouse *(Centrocercus urophasianus*, hereafter sage-grouse) is a large sexually dimorphic tetraonid that is endemic to the sagebrush *(Artemisia* spp.) biome of western North America (Knick and Connelly, 2011). Each spring, adult males aggregate in open areas called leks where they display for females. Fertilized females then nest on the ground among the sagebrush (Holloran and Anderson, 2005). Initially, the chicks feed on insects before switching to forbs. The adults predominantly feed on sagebrush, especially in the winter. Most males begin lekking two years after hatching. Mean peak counts of males on leks are commonly used as an abundance metric (Connelly and Braun, 1997; Doherty et al., 2010; Fedy and Aldridge, 2011)

A multitude of studies have reported local negative effects of oil and gas (OAG) development on sage-grouse lek counts, movement, stress-levels and fitness components. The most frequently-reported phenomenon is the decline in lek counts with increasing densities of well pads (Walker et al., 2007; Doherty et al., 2010; Harju et al., 2010; Green et al., 2016). Reductions in fitness components such as lower nest initiation rates (Lyon and Anderson, 2003) and lower annual survival of yearlings reared in areas where OAG infrastructure is present (Holloran et al., 2010) have been detected using radio-tracking. The development of Global Positioning System (GPS) telemetry methods has facilitated the fitting of more sophisticated and realistic spatially-explicit habitat use models which suggest that nest and brood failure is influenced by proximity to anthropogenic features (Dzialak et al., 2011). More recently, experimental studies have suggested that noise alone can reduce lek attendance (Blickley et al., 2012b) and increase stress hormones (Blickley et al., 2012a).

However, to date no-one has examined whether sage-grouse population-level responses are consistent with the local studies. Although Green et al. (2016) state that they modeled sage-grouse populations, they use their population dynamic models to analyze the effects of OAG on changes in abundance at individual leks. Even authors such as Walker et al. (2007) and Gamo and Beck (2017) who analyzed aggregations of leks, group their leks by level of OAG development as opposed to population boundaries.

Although it has received less attention than OAG, climatic variation has also been shown to influence sage-grouse lek counts, survival, clutch size and nesting success (Blomberg et al., 2012, 2014, 2017; Coates et al., 2016; Gibson et al., 2017). This is not surprising, as there is a long and ecologically important history of studies on the influence of climatic variation on the population dynamics of tetraonids (Moran, 1952, 1954; Ranta et al., 1995; Lindström et al., 1996; Cattadori et al., 2005; Ludwig et al., 2006; Kvasnes et al., 2010; Selås et al., 2011; Viterbi et al., 2015; Ross et al., 2016). Consequently, the current study also includes annual variation in regional climate as a potential predictor of sage-grouse population dynamics.

Previous studies of the effect of climatic variation on sage-grouse have used local temperature and precipitation data with mixed results (Blomberg et al., 2012; Green et al., 2016; Blomberg et al., 2014, 2017; Coates et al., 2016; Gibson et al., 2017; Green et al., 2016). However, large-scale climate indices often outperform local data in predicting population dynamics and ecological process (Stenseth et al., 2002; Hallett et al., 2004). The Pacific Decadal Oscillation (PDO), which is derived from the large-scale spatial pattern of sea surface temperature in the North Pacific Ocean (Mantua et al., 1997), is potentially the most important climatic process influencing the sagebrush biome (Neilson et al., 2005). Consequently, the PDO index was chosen as the climate indicator.

Wyoming was selected for the current study because it contains approximately 37% of the recent range-wide population of sage-grouse (Copeland et al., 2009; Fedy et al., 2012), has experienced substantial levels of OAG development dating to the late 1800s (Braun et al., 2002) and because the lek location and count data were available for research.

## METHODS

### Data Preparation

#### Sage-grouse Data

When multiple counts exist for the same lek in a single year, almost all authors take the maximum count (Holloran, 2005; Walker et al., 2007; Harju et al., 2010; Fedy and Aldridge, 2011; Fedy and Doherty, 2011; Garton et al., 2011; Blickley et al., 2012b; Blomberg et al., 2013; Davis et al., 2014; Garton et al., 2015; Coates et al., 2016; Fremgen et al., 2016; Monroe et al., 2016; Green et al., 2016). The justification for using the maximum count is articulated by Garton et al. (2011) who state that,

> …counts over the course of a single breeding season vary from a low at the beginning of the season, to peak in the middle, followed by a decline to the end, which necessitates using the maximum count from multiple counts across the entire season as the index.

However, as noted by Johnson and Rowland (2007), this results in a substantial upward bias at leks with multiple counts. To understand why consider an unbiased die. The expectation with a single throw is 3.5. With two throws the expectation for the mean value is still 3.5 but the expectation for the maximum value is 4.47. To avoid this bias, several alternative approaches are available: exclude early and late counts and then either include the repeated counts in the model (Gregory and Beck, 2014) or take the mean of the repeated counts and/or explicitly model the change in attendance through time (Walsh et al., 2004) as is done for spawning salmon (Hilborn et al., 1999). We excluded early and late counts and took the rounded mean of the repeated counts. However as discussed below we also assessed the sensitivity of the results to the use of the rounded mean as opposed to maximum count.

The sage-grouse lek count and location data were provided by the State of Wyoming. After excluding male lek counts with unknown counts or dates or those before 1985 there were 88,771 records. To reduce potential biases, only the most reliable male lek counts were included in the analyses. In particular, only ground counts from leks that were checked for activity and were part of a survey or count were included (as per Wyoming Game and Fish guidelines). This reduced the number of records to 79,857. To ensure counts were close to the peak (see above), only data that were collected between April 1st and May 7th were included. This reduced the number of records to 65,439. Finally, lek counts for which the number of individuals of unknown sex were ≥ 5% of the number of males (suggesting unreliable identification) were excluded which left a total of 42,883 records. The leks with at least one remaining count are mapped in Figure 1 and the associated mean male lek counts are plotted in Figure 2.

**Figure 1.**
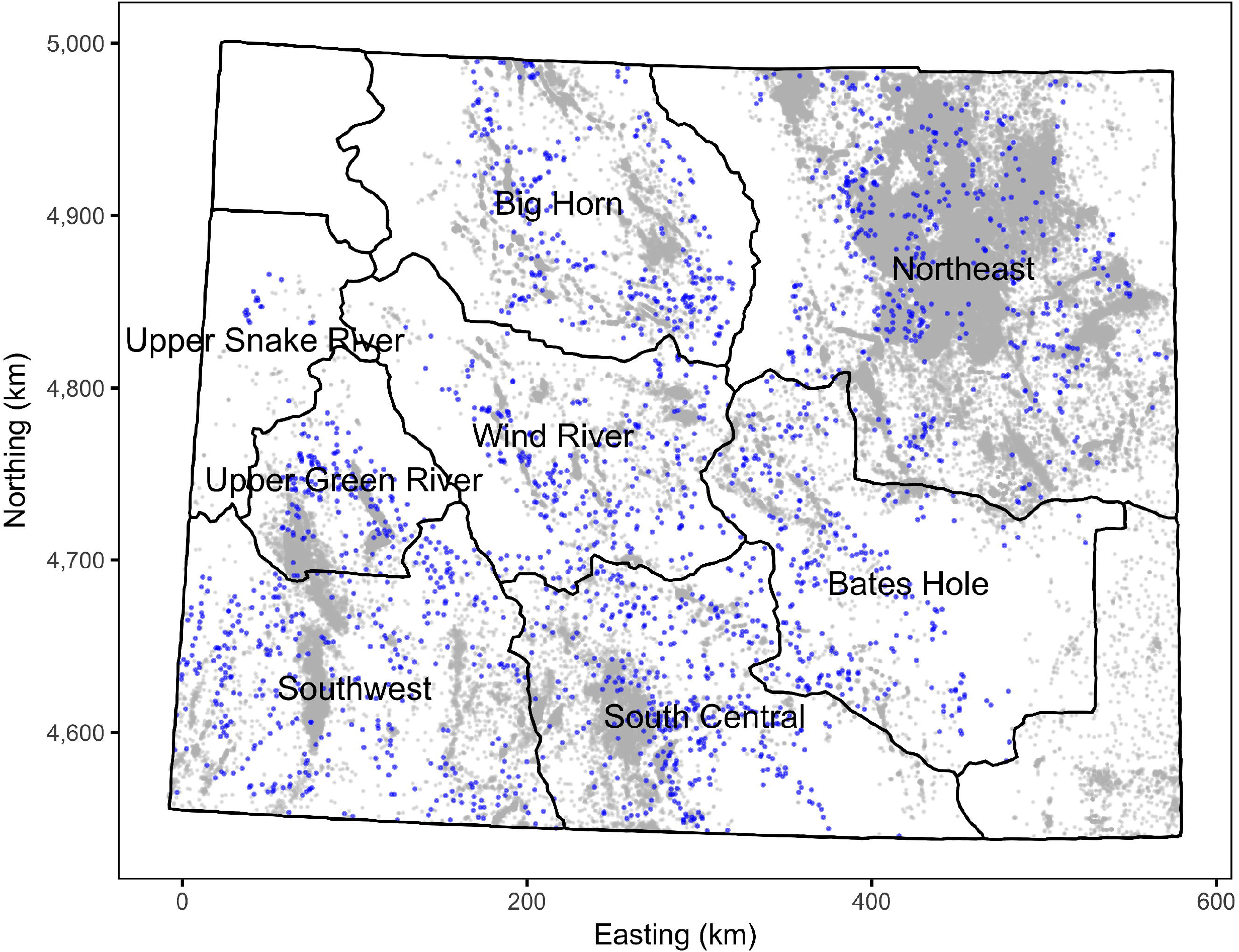
Map of Wyoming and its working groups. Leks are indicated by blue points and well pads by grey points. Only leks and wells pads that are included in the analyses are shown. The leks and well pads are not to scale. The projection is EPSG:26913.

**Figure 2.**
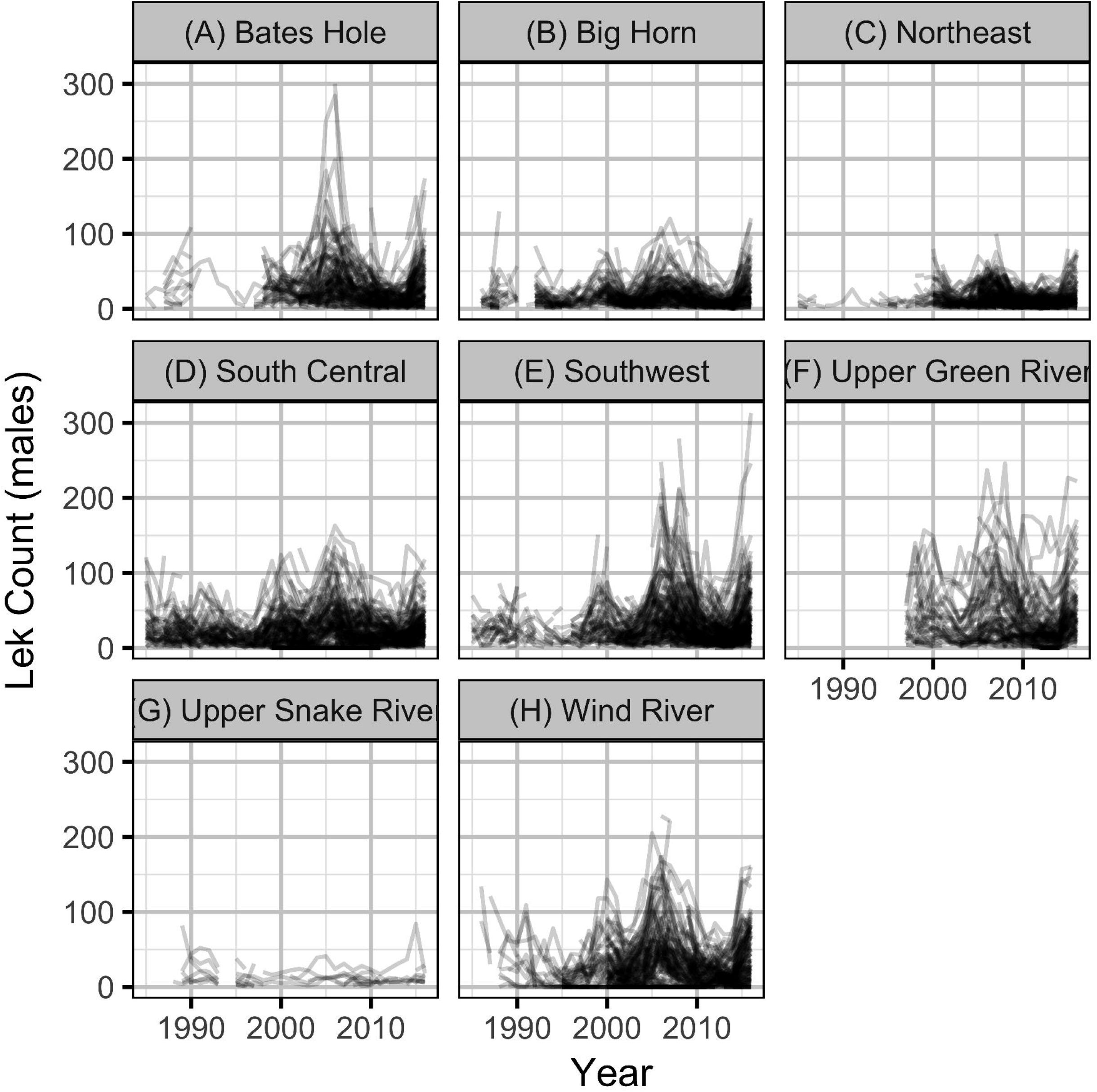
Mean counts of male sage-grouse at individual leks by year and working group.

The State of Wyoming utilizes eight regional sage-grouse working groups to facilitate local population management and data reporting, including lek counts and hunting harvest (Fig. 1, Christiansen and Belton, 2017). For the purposes of the current study, we also treat these working groups as if they are separate populations. The population densities (males per lek) were calculated by averaging the mean counts for individual leks for each working group in each year.

#### Oil and Gas Data

Wyoming Oil and Gas Conservation Commission (WOGCC) conventional, coal-bed and injection well pad location and production data were downloaded from the Wyoming Geospatial Hub (http://pathfinder.geospatialhub.org/datasets/) at 2018-05-25 02:13 UTC. Well pads without a provided spud date were excluded as were well pads constructed before 1900 or after 2016. The included well pads are mapped in Figure 1.

The intensity of OAG development was quantified in terms of the proportional areal disturbance due to well pads within a specific distance of the leks. The areal disturbance was calculated at lek distances of 0.8, 1.6, 3.2 and 6.4 km with the areal disturbance of each well pad considered to have a radius of 60 m (Green et al., 2016). The annual areal disturbances for individual leks with lek counts at 3.2 km are plotted in Figure 3.

**Figure 3.**
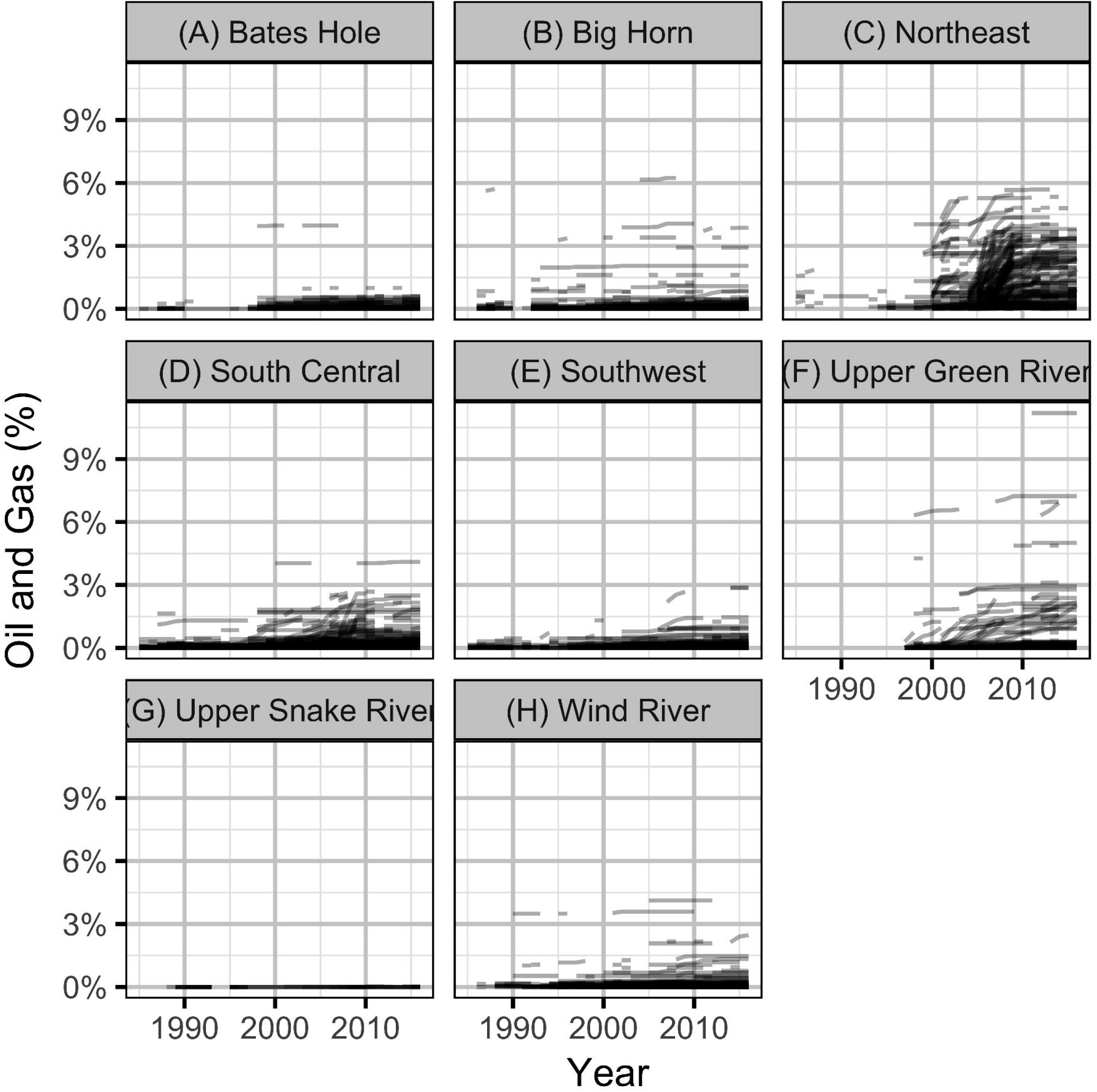
Percent areal disturbance due to well pads within 3.2 km of individual leks with one or more counts by year and working group.

#### Climatic Data

The PDO index (Trenberth and Hurrell, 1994; Mantua et al., 1997) data were queried from the rpdo R package (Fig. 4).

**Figure 4.**
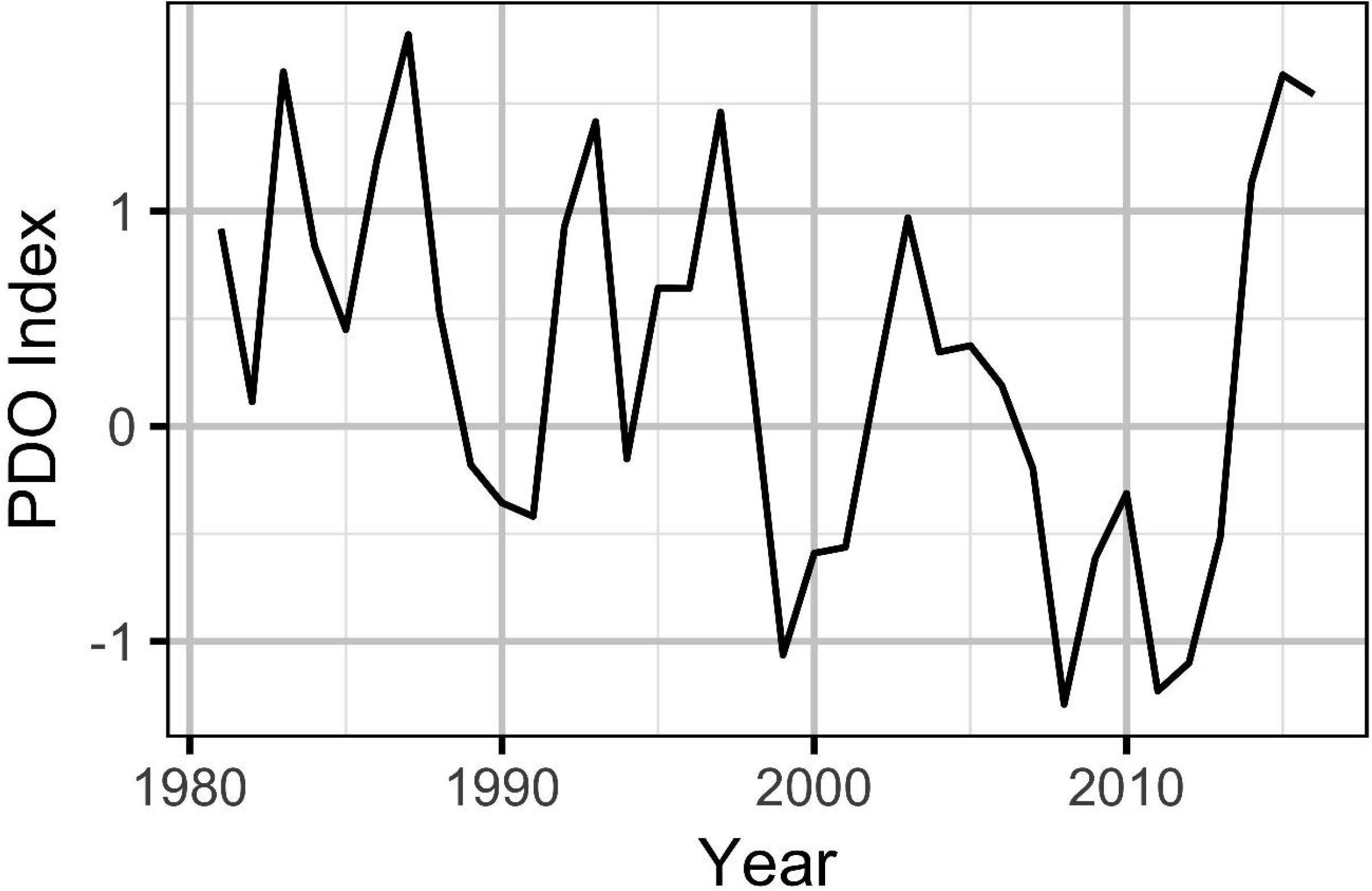
Pacific Decadal Oscillation index by year. Positive values indicate a warm phase and negative values a cool phase.

### Statistical Analysis

#### Local Models

The individual lek counts were analyzed using GLMMs (Bolker et al., 2009) with the standardized areal disturbance due to OAG and the PDO index as fixed effects and year and lek as random effects. The areal disturbance and PDO index were standardized (centered and divided by the standard deviation) to facilitate comparison. As preliminary analysis indicated that the lek counts were overdispersed, the GLMMs utilized a negative binomial distribution.

More formally, the lek count model is described by the following equations

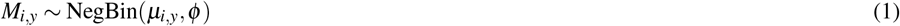

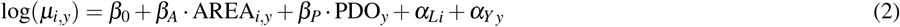

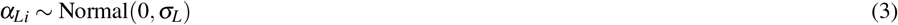

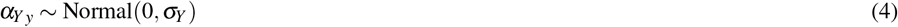

where *M_i,y_* is the rounded mean count of males for the ith lek in the *y*th year, *β_A_* and *β_P_* are the fixed effects of the standardized areal disturbance due to well pads (AREA_*i,y*_) and PDO index (PDO_*y*_) on the expected count (*μ_i,y_*), *σ_L_* and *σ_Y_* are the standard deviations (SDs) of the random effects of lek and year. In our parameterization of the negative binomial the parameter *φ* controls the overdispersion scaled by the square of *μ*, i.e.,

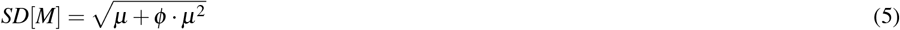

Key model parameters are also described in Table 1.

**Table 1.** Descriptions of key model parameters.

| Parameter | Description |
| --- | --- |
| $\beta_0$ | The intercept for the log lek count or log population density. |
| $\beta_D$ | The effect of population density on $\beta_0$. |
| $\beta_P$ | The effect of the standardised Pacific Decadal Oscillation index on $\beta_0$. |
| $\beta_A$ | The effect of the standardised areal disturbance due to well pads on $\beta_0$. |
| $\phi$ | The overdispersion term. |
| $\sigma_\varepsilon$ | The SD of the observer error. |
| $\beta_N$ | The intercept for the SD of the process error. |
| $\beta_L$ | The effect of the log of the number of leks counted on $\beta_N$. |
| $\sigma_G$ | The SD of the random effect of working group on $\beta_0$. |
| $\sigma_L$ | The SD of the random effect of lek on $\beta_0$. |
| $\sigma_Y$ | The SD of the random effect of year on $\beta_0$. |

To identify the most important spatial scale (distance from each lek when calculating the areal disturbance) and temporal lags, a total of 64 models were fitted to the lek count data representing all combinations of the four lek distances (0.8, 1.6, 3.2 and 6.4 km) and independent lags of one to four years in the areal disturbance (Walker et al., 2007; Doherty et al., 2010; Harju et al., 2010; Gregory and Beck, 2014) and PDO index. The relative importance of each spatial scale and temporal lag as a predictor of individual lek counts was assessed by calculating it’s Akaike’s weight (*w_i_*) across all 64 models (Burnham and Anderson, 2002).

Once the model with the most important spatial scale and temporal lags was identified, the relative importance of *β_A_* and *β_P_* was quantified by calculating their Akaike’s weights across the selected full model and the three reduced variants representing all combinations of the two parameters (Burnham and Anderson, 2002) and by calculating their effect sizes with 95% confidence/credible limits (CLs Bradford et al., 2005; Claridge-Chang and Assam, 2016). The effect sizes, which represent the expected percent change in the lek count with an increase in the predictor of one SD, were calculated for the final full model and by averaging across all four models (Burnham and Anderson, 2002; Turek, 2015).

#### Population Models

The calculated annual population densities (mean males per lek) in each working group were analyzed using Gompertz Population Dynamic Models (Garton et al., 2011) with the standardized areal disturbance and PDO index as fixed effects and year and group as random effects. Gompertz Population Dynamic Models (GPDMs) were used because they incorporate density-dependence (Dennis et al., 2006; Knape and de Valpine, 2012) and have performed well in explaining rates of change for sage-grouse in general and for Wyoming sage-grouse in particular (Garton et al., 2011).

The population model is described by the following equations

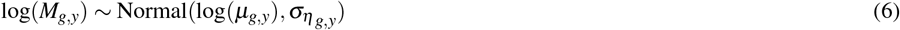

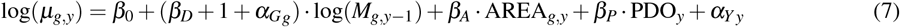

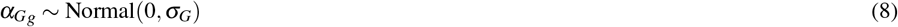

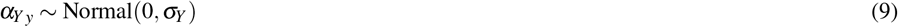

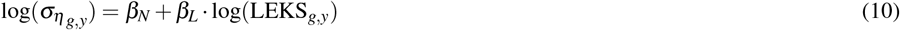

where *M_g,y_* is the density at the gth group in the yth year, *μ_g,y_* is the expected density, *β_D_* is the typical density-dependence and *α_G_g__* is the group-level random effect on the density-dependence, *σ_η_g,y__* is the expected process error (Dennis et al., 2006), *β_N_* is the intercept for the log process error and *β_L_* is the effect of the number of leks surveyed on *β_N_*. The other terms are approximately equivalent to those in the lek count model. The equivalence is only approximate as the terms in the population model act on the change in density (as opposed to density).

The carrying capacity, which represents the long-term expected density around which a population fluctuates (Dennis et al., 2006), is given by

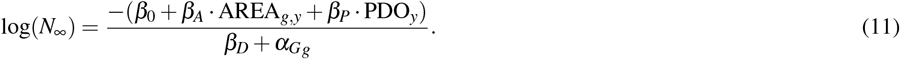

Preliminary analyses considered Gompertz State-Space Population Models (Dennis et al., 2006; Knape and de Valpine, 2012) which estimate both process and observer error (Maunder et al., 2015). However, the models were unable to reliably estimate both error terms. As the observer error was estimated to be smaller than the process error and because ignoring process error can bias Akaike’s Information Criterion based tests towards incorrectly accepting covariates (Maunder et al., 2015), we followed Garton et al. (2011) in assuming no observer error. The preliminary analyses indicated that fixing the observer error at zero had little effect on the results. The process error was allowed to vary by the number of leks surveyed, which varied from one to 214, to attempt to account for the additional stochasticity associated with smaller populations and/or lower coverage.

The primary question this study attempts to answer is whether the sage-grouse population-level responses to oil and gas are consistent with the local studies. Consequently, the average areal disturbance in each working group was calculated at the spatial scale that was most important in the local analyses. However, as the timing of effects could differ between the local count models and the population dynamic models, the Akaike’s weight for each lag of one to four years in the areal disturbance and one to four years in the PDO index was calculated across the 16 models representing all lag combinations. Once the full model with the most important temporal lags had been identified, the relative importance of *β_A_* and *β_P_* was once again quantified from their effect sizes with 95% CLs and their Akaike’s weights across the full model and the three reduced variants.

To assess the sensitivity of the population model outputs to the inputs, a local, qualititative sensitivity analysis (Pianosi et al., 2016) was conducted. More specifically, the effect sizes in the full model were estimated using 1) the maximum of the repeated counts (Garton et al., 2011) as opposed to the mean (Johnson and Rowland, 2007) and 2) the data from 1997 and 2005 onwards. The former period was used during preliminary analyses due to the availability of hunter-harvested wing count data (Braun and Schroeder, 2015). From 2005 onwards regulatory and technological developments were introduced to reduce the impacts of OAG (Applegate and Owens, 2014) on sage-grouse. The effect sizes were adjusted for any differences in the SDs of the data.

### Predicted Population Impacts

To examine whether the population-level responses are consistent with the lek-level results, the expected effect of OAG on the long-term mean densities in each working group was calculated for the local and population models. In the case of the local model, the predicted population impacts represent the percent difference in the sum of the expected counts for the observed levels of OAG versus no OAG across all leks in the working group after accounting for annual and climatic effects. In the case of the population model, the predicted population impacts represent the percent difference in the expected carrying capacity with the observed levels of OAG versus no OAG accounting for annual and climatic effects.

### Statistical Methods

For reasons of computational efficiency, the initial 64 local and 16 population-level models were fit using the frequentist method of Maximum Likelihood (ML, Millar, 2011). The Akaike’s weights were calculated from the marginal Akaike’s Information Criterion values corrected for small sample size (mAICc, Burnham and Anderson, 2002; Vaida and Blanchard, 2005; Greven and Kneib, 2010). Model adequacy was assessed by plotting and analysis of the standardized residuals from the final full ML model (Burnham and Anderson, 2002) with the most important spatial scale and lags. As both the local and population models used log-link functions, the effect sizes (percent change in the response for an increase in one SD) were calculated from exp(*β*) – 1 where *β* is the fixed effect of interest or its upper or lower CL. The ML effect sizes were calculated from the full model and averaged across the full model and three reduced variants (Lukacs et al., 2010).

To allow the predicted population impacts to be estimated with CLs, the final full models were also fitted using Bayesian methods (Gelman et al., 2014). The prior for all primary parameters was an uninformative (Gelman et al., 2014) normal distribution with a mean of 0 and a SD of 5. A total of 1,500 MCMC samples were drawn from the second halves of three chains. Convergence was confirmed by ensuring that Rhat was ≤ 1.01 (Gelman et al., 2014) and the effective sample size was ≥ 1,000 (Brooks and Gelman, 1998) for each structural parameter.

### Software

The data preparation, analysis and plotting were performed using R version 3.5.0 (R Core Team, 2017) and the R packages TMB (Kristensen et al., 2016) and rstan (Stan Development Team, 2016). The clean and tidy analysis data and R scripts are archived at https://doi.org/10.5281/zenodo.837866. The raw sage-grouse data, which provide the lek locational information, are available from the Wyoming Department of Fish and Game. The raw data are not required to replicate the analyses.

## RESULTS

### Local Models

The Akaike weights for the spatial scales indicate that 3.2 km is unanimously supported (*w_i_* = 1.00) as the most important lek distance for predicting individual lek counts from the areal disturbance due to well pads (Table S1). The Akaike weights for the lags in the areal disturbance provided close to unanimous support for a single candidate with the lag of one year receiving a weight of 0.99 (Table S2). The situation with the PDO index lags was less clear-cut (Table 2), although a lag of two years received the majority of the support (*w_i_* = 0.73). Consequently, the local model with a lek distance of 3.2 km and lags of one and two years in the areal disturbance due to well pads and the PDO index, respectively, was selected as the final model. The standardized residuals, with the exception of a small number of high outliers, were approximately normally distributed and displayed homogeneity of variance. Most leks had an expected count of male sage-grouse in the absence of OAG of approximately 10 birds (Fig. S1).

**Table 2.** The relative importance (*w_i_*) of the lag in the Pacific Decadal Oscillation index density as a predictor of the count of males sage-grouse at individual leks across all models with a lek distance of 0.8, 1.6, 3.2, 6.4 and 12.8 km and the areal disturbance due to well pads independently lagged one to four years.

| PDO Lag (yr) | Models | Proportion | $w_i$ |
| --- | --- | --- | --- |
| 2 | 16 | 0.25 | 0.73 |
| 3 | 16 | 0.25 | 0.18 |
| 4 | 16 | 0.25 | 0.05 |
| 1 | 16 | 0.25 | 0.04 |

The Akaike weights for *β_A_* (*w_i_* = 1) and *β_P_* (*w_i_* = 0.98) across the final full model and the three reduced models indicate that both are very strongly supported as predictors of individual lek counts. The effect size estimates (Fig. 5) indicate that OAG and the PDO have large negative (Fig. 6) and positive (Fig. 7) impacts of similar magnitudes (just under 20%) on the lek counts and that the estimates are insensitive to the statistical framework (ML of Bayesian) or model-averaging (Tables S3-S5). Despite the inclusion of the PDO index as an important predictor, there was still substantial remaining annual cyclical variation in the lek counts (Fig. S2) which was modeled by the random effect of year.

**Figure 5.**
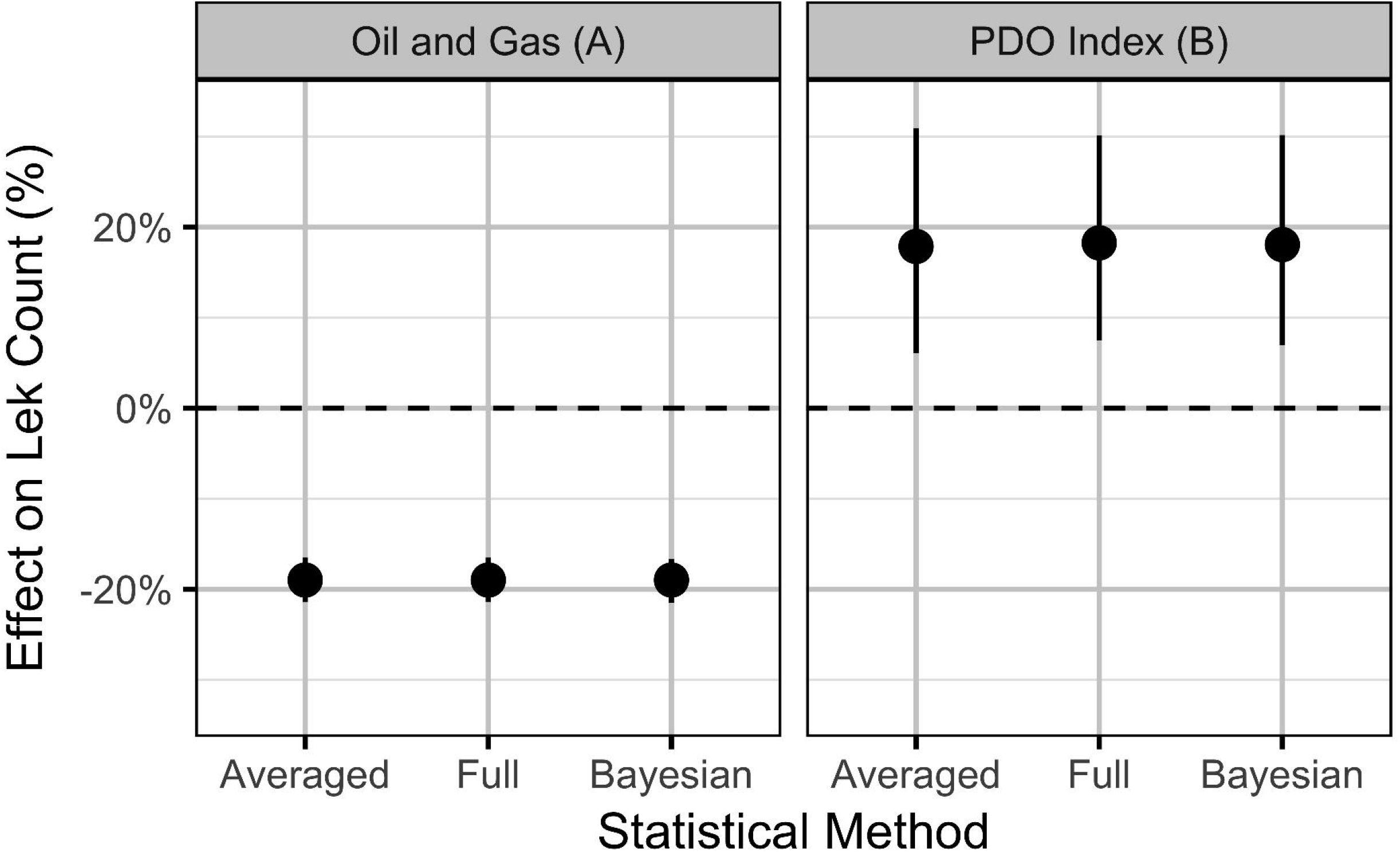
Estimates (with 95% CIs) of the effect of an increase in one SD (0.8 %) in the areal disturbance due to well pads within 3.2 km and the Pacific Decadal Oscillation index (0.82). The effect is on the expected count of male sage-grouse at an individual lek.

**Figure 6.**
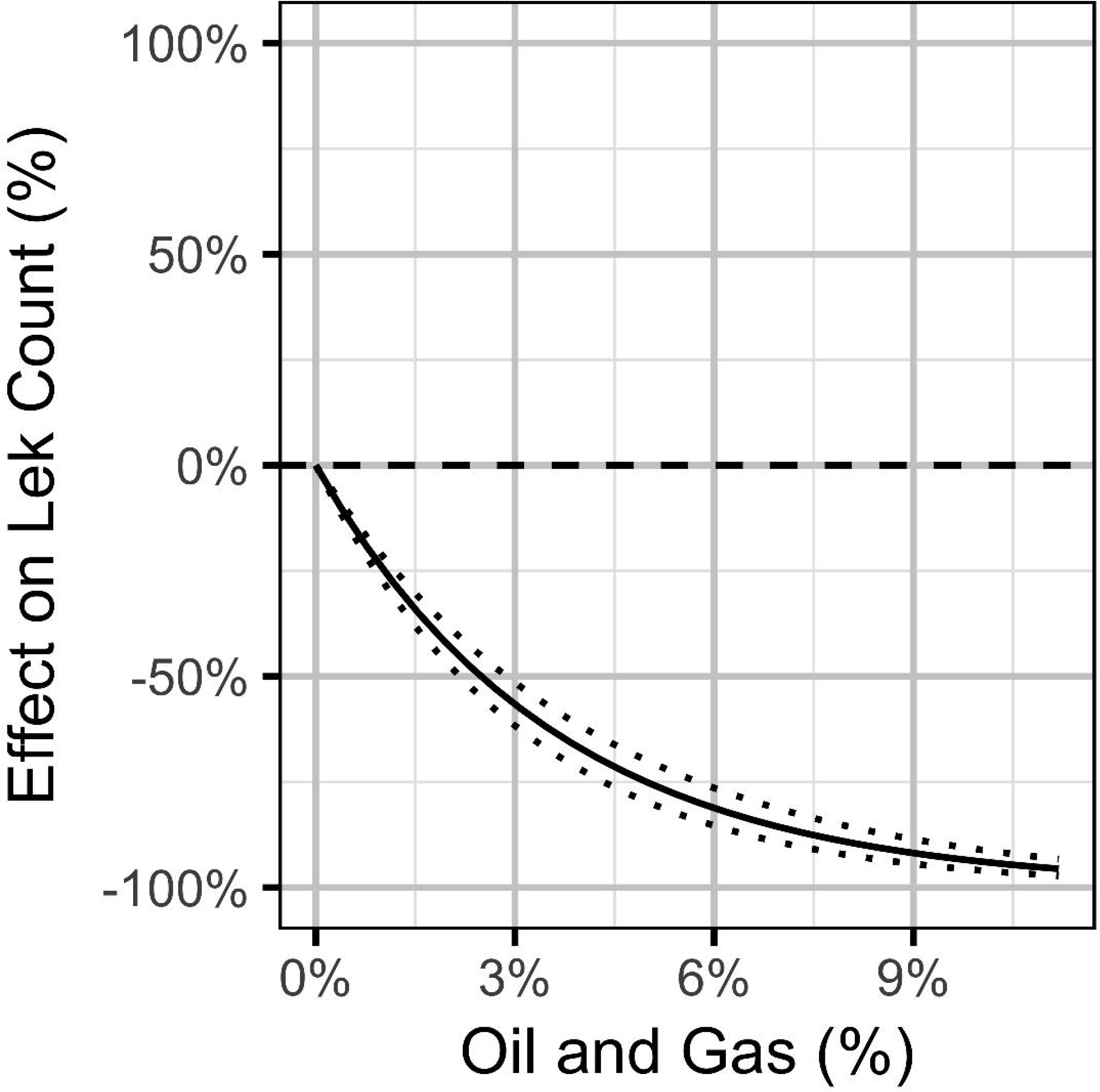
Bayesian estimates (with 95% CIs) of the effect of the percent areal disturbance due to oil and gas well pads on the expected count of male sage-grouse at a typical lek. The effect is the percent change in the expected count relative to no areal disturbance.

**Figure 7.**
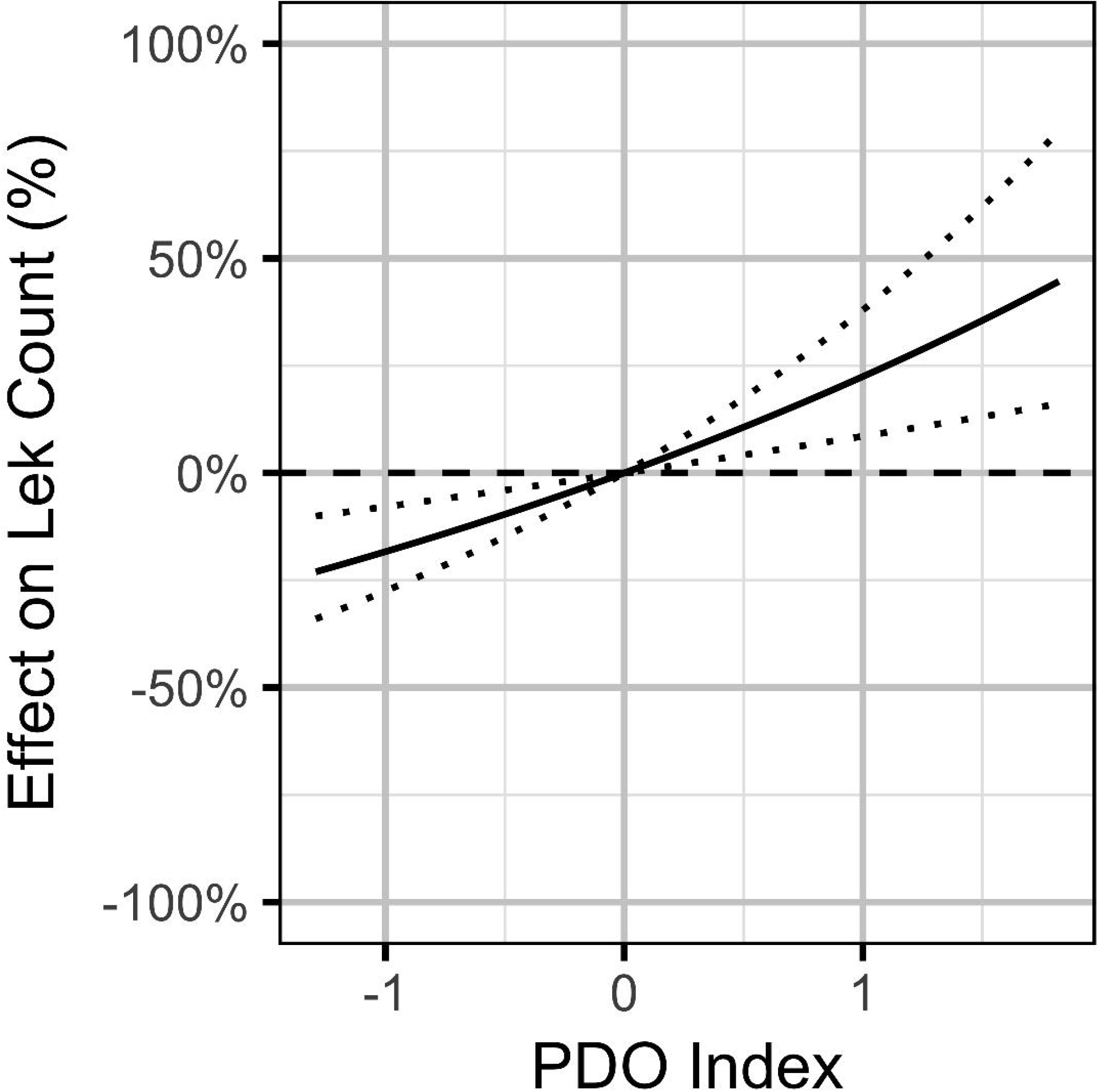
Bayesian estimates (with 95% CIs) of the effect of the Pacific Decadal Oscillation index on the expected count of male sage-grouse at a typical lek. The effect is the percent change in the expected count relative to a Pacific Decadal Oscillation index value of 0.

### Population Models

Based on the results of the local models, the level of OAG development in each working group was calculated in terms of the average areal disturbance due to well pads within 3.2 km of each lek (Fig. 8). The Akaike weights for the lag in the areal disturbance (Table 3) were largely indifferent (0.29 to 0.20) although a lag of one year had the most support. The Akaike weights for the PDO index (Table 4) provided the majority of the support for a lag of one year (*w_i_* = 0.53). The model predictions provided a reasonable fit to the annual mean lek counts which exhibit large cyclical fluctuations (Fig. 9). The residuals were approximately normally distributed with homogeneity of variance. The carrying capacities in the absence of OAG varied between approximately 13 males per lek in the Upper Snake River to approximately 35 males per lek in the Upper Green River (Fig. S3).

**Figure 8.**
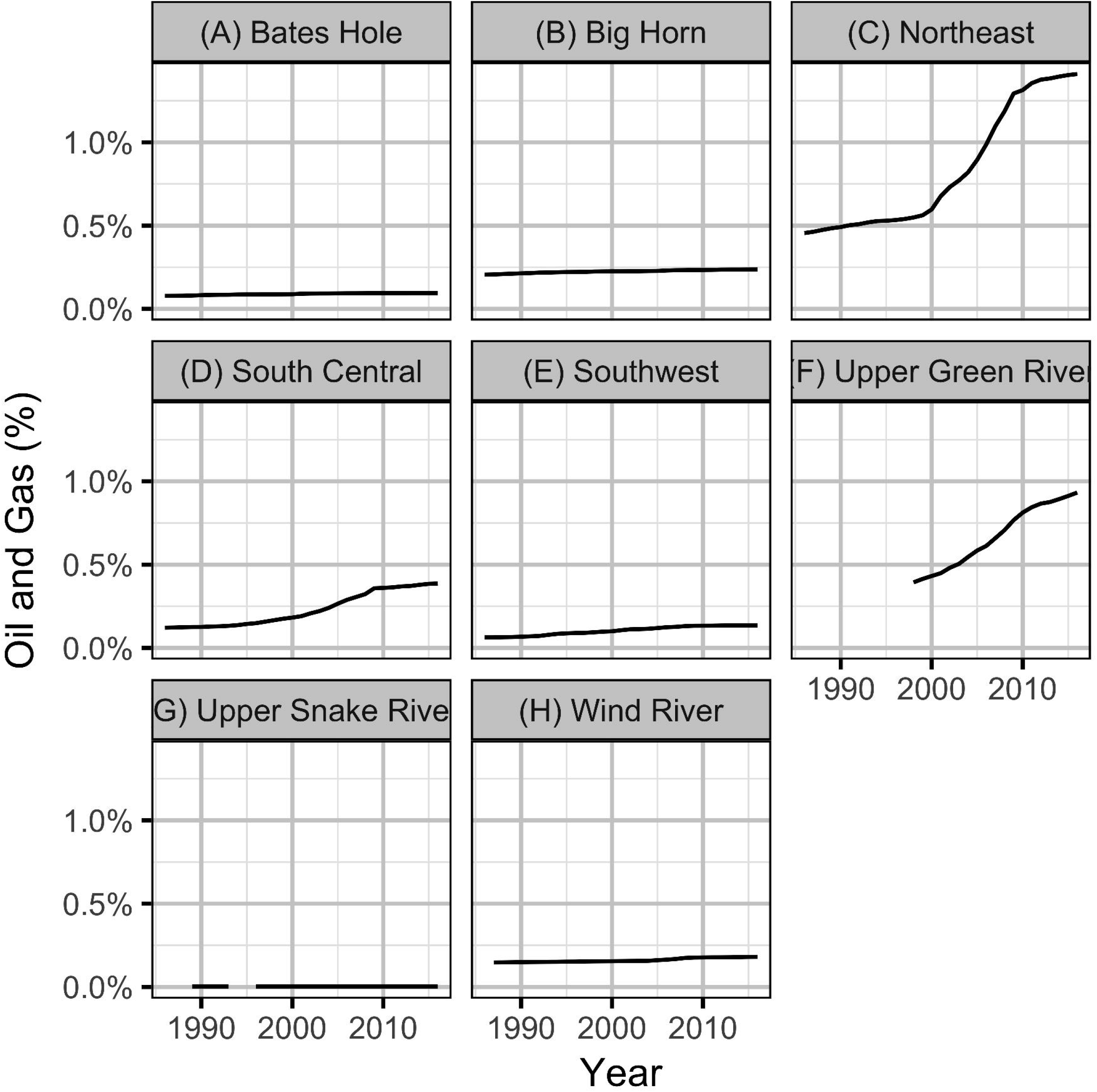
Mean areal disturbances due to well pads within 3.2 km of all leks by year and working group.

**Figure 9.**
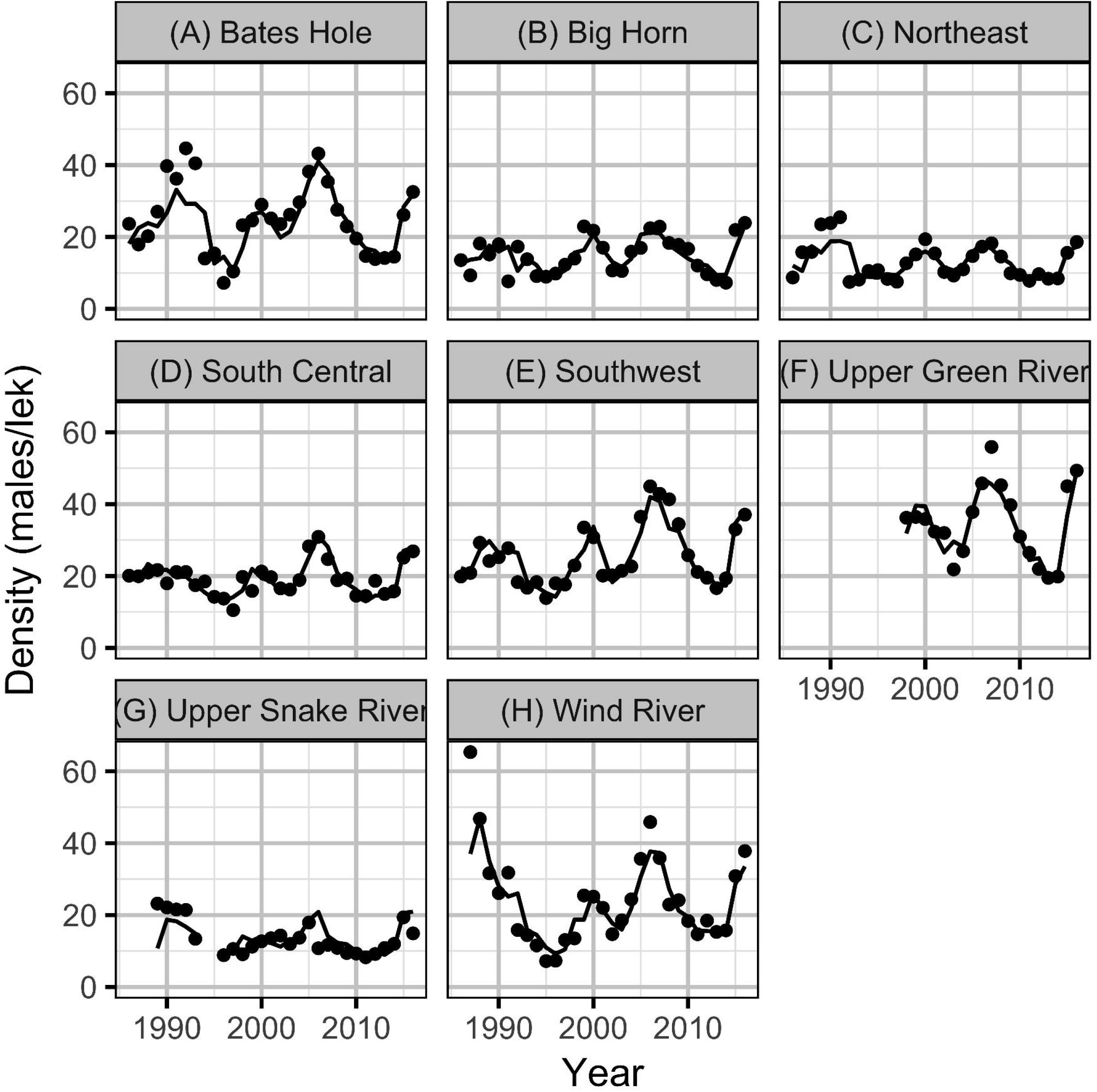
Mean lek counts by year and working group. The solid line is the estimate of the population density based on the observed density in the previous year for the final full Maximum Likelihood model.

**Table 3.** The relative importance (*w_i_*) of the lag in the areal disturbance due to well pads as a predictor of the change in the population density across all models with a lek distance of 3.2 km and the Pacific Decadal Oscillation index independently lagged one to four years.

| Area Lag (yr) | Models | Proportion | $w_i$ |
| --- | --- | --- | --- |
| 1 | 4 | 0.25 | 0.29 |
| 2 | 4 | 0.25 | 0.27 |
| 3 | 4 | 0.25 | 0.24 |
| 4 | 4 | 0.25 | 0.20 |

**Table 4.** The relative importance (*w_i_*) of the lag in the Pacific Decadal Oscillation index as a predictor of the change in the population density across all models with a lek distance of 3.2 km and the areal disturbance due to well pads lagged one to four years.

| PDO Lag (yr) | Models | Proportion | $w_i$ |
| --- | --- | --- | --- |
| 1 | 4 | 0.25 | 0.53 |
| 2 | 4 | 0.25 | 0.32 |
| 3 | 4 | 0.25 | 0.08 |
| 4 | 4 | 0.25 | 0.08 |

The Akaike weights for *β_A_* and *β_P_* (Table 1) across the final full model and the three reduced models indicate that while the PDO index receives moderate support (*w_i_* = 0.74) as a predictor of population changes, there is only weak support (*w_i_* = 0.41) for the areal disturbance. In addition, the effect size estimates (Fig. 10), which are sensitive to model-averaging but not the statistical framework (Tables S6-S8), indicate that while the PDO has a moderate positive influence (effect size of approximately 8%) the effect of OAG on the subsequent year’s density is relatively small (effect size of −2.5%, Fig. 10). However, despite it’s relatively small effect size, OAG may have a substantial effect (−20% reduction) on the long-term carrying capacity in the most impacted working groups (Fig. 11). The reasons why are discussed below. The effect of the PDO on the carrying capacity (Fig. S4) is comparable to its effect on the counts at individual leks although there is much more uncertainty (Fig. 7). As for the local models, the random effect of year accounted for substantial unexplained annual cyclical variation (Fig. S5). The effect of density on the subsequent year’s density (i.e.density-dependence) is relatively minor: in a typical working group a reduction in the density to 10 males (half the carrying capacity) results in an average of just 13 males the following year (Fig. S6).

**Figure 10.**
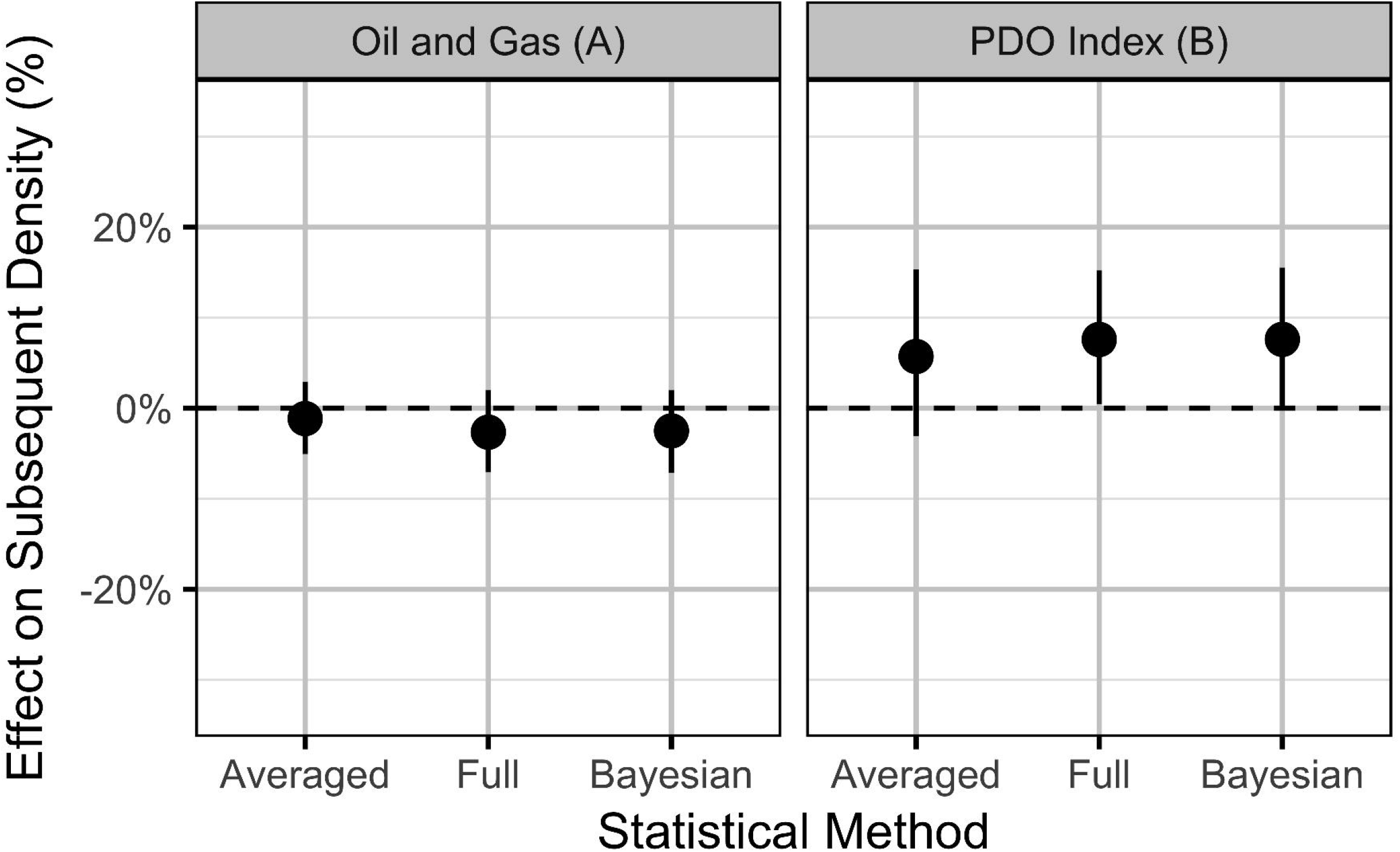
Estimates (with 95% CIs) of the effect of an increase in one SD (0.31%) in the areal disturbance due to well pads within 3.2 km of all leks and the Pacific Decadal Oscillation index (0.86). The effect is on the expected subsequent population density.

**Figure 11.**
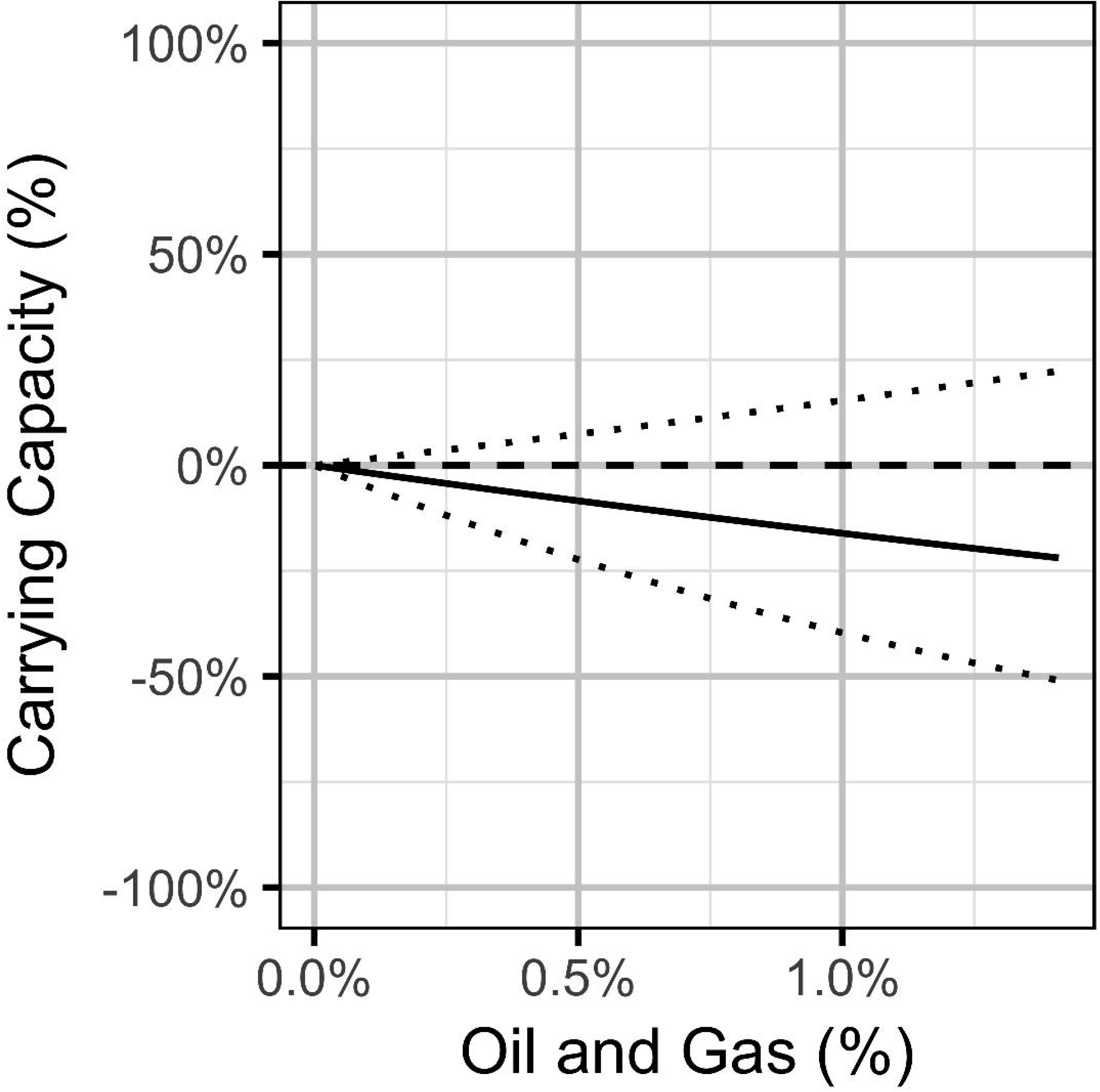
Bayesian estimates (with 95% CIs) of the effect of the percent areal disturbance due to oil and gas well pads on the expected carrying capacity at a typical working group. The effect is the percent change in the expected carrying capacity relative to no areal disturbance.

The qualititative sensitivity analysis indicates that using the maximum as opposed to the rounded mean of the repeated lek counts has a negligible effect on the effect size estimates (Fig. 12). In contrast, the estimated effect of the PDO is strongest in the most recent data.

**Figure 12.**
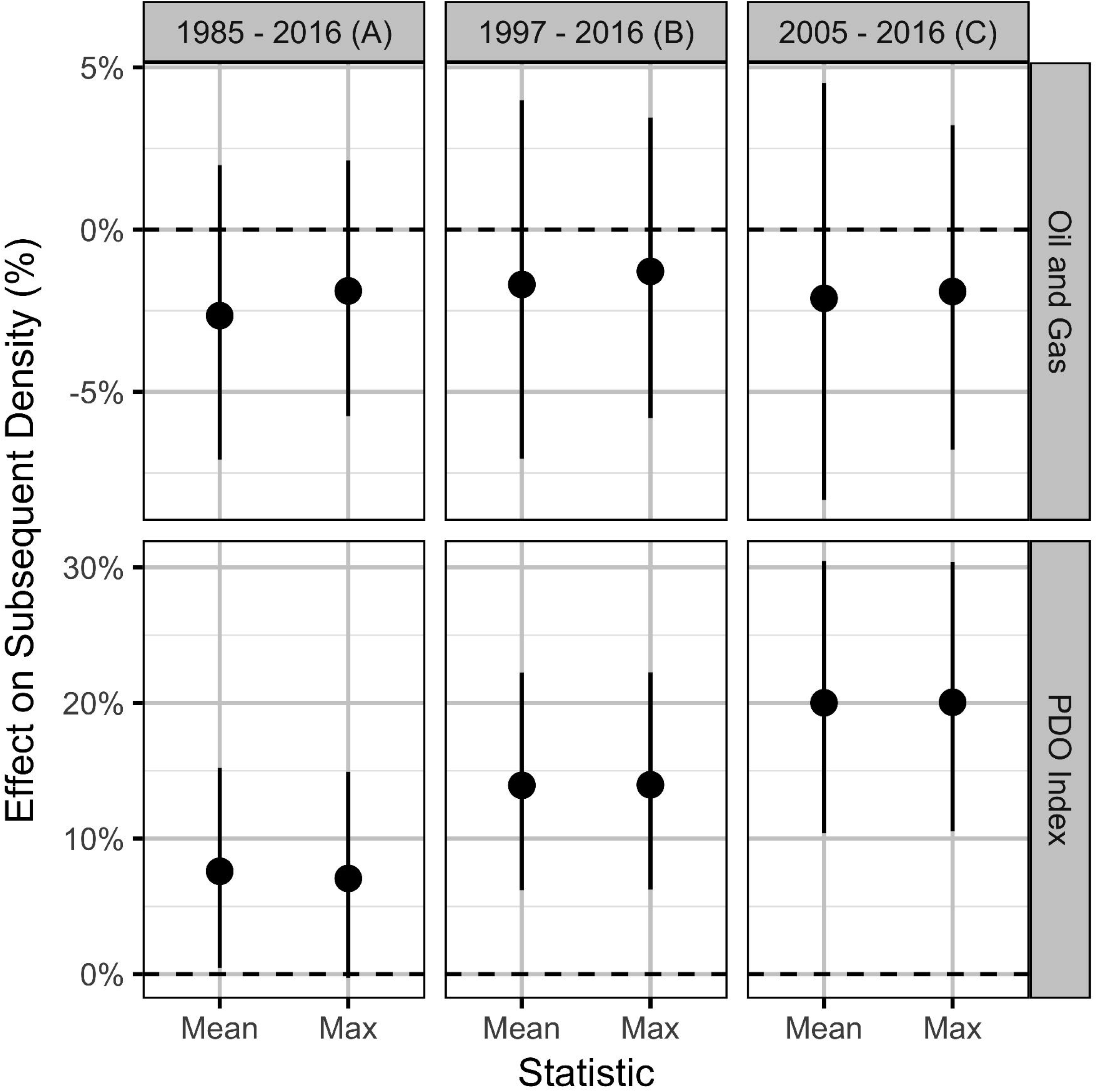
Estimates (with 95% CIs) of the effect of an increase in one SD (0.31%) in the areal disturbance due to well pads within 3.2 km of all leks and the Pacific Decadal Oscillation index (0.86) by the statistic used to combine repeated lek counts and the time period. The effect is on the expected subsequent population density. When the time interval is limited to data from more recent years where a greater number of leks were surveyed and more frequently (i.e. data are of higher quality) the effect of oil and gas becomes less certain, while the PDO has a greater effect.

### Predicted Population Impacts

The predicted population impacts from the Bayesian full models indicate that although OAG has a relatively minor influence on the change in the population density, the possibility of a larger negative effect on the carrying capacity that is consistent with summation of the local lek-level impact could not be excluded (Fig. 13).

**Figure 13.**
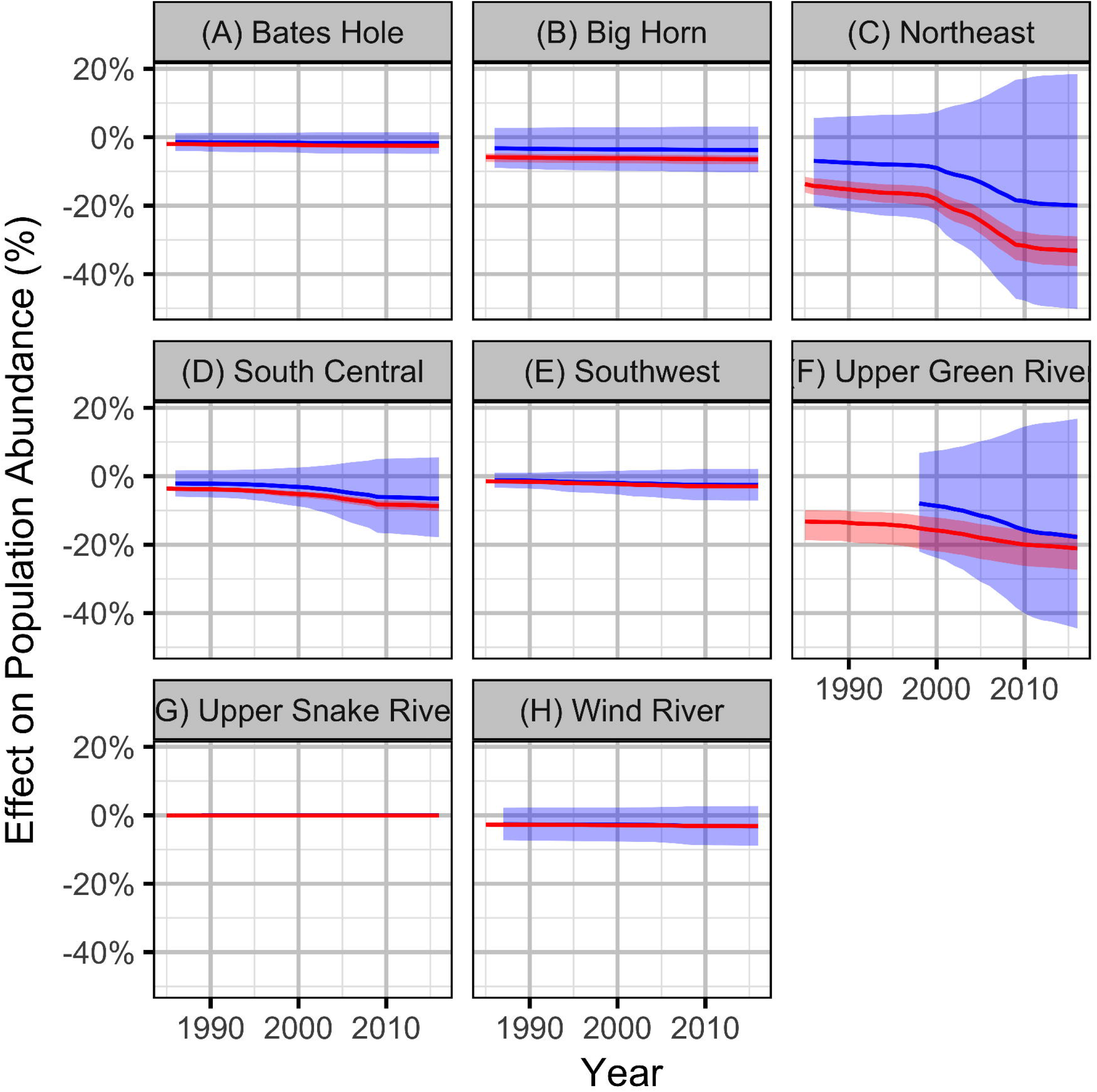
Bayesian estimates (with 95% CIs) of the effect of the observed levels of oil and gas on the population abundance of sage-grouse based on the local (red) and population (blue) models.

## DISCUSSION

### Oil and Gas

This is not the first study to consider whether local impacts potentially extend to population declines. Based on local declines, Copeland et al. (2013) estimated that sage-grouse populations in Wyoming will decrease by 14 to 29%, but that a conservation strategy that includes the protection of core areas could reduce the loss to between 9 and 15% while Copeland et al. (2009) estimated that future OAG development in the western United States (US) will cause a long-term 7 to 19% decline in sage-grouse numbers relative to 2007. As argued by Doherty et al. (2010), estimation of population-level impacts is important because it provides a biologically-based currency for quantifying the cost of OAG as well as the benefits of mitigation or conservation.

This is however the first study to examine whether the actual population-level response is consistent with the local impacts. A key conclusion is that scaling up the local, lek level impacts produces estimates of population declines similar to those for the individual populations although there is much uncertainty over the magnitude, if any, of the actual population-level response.

Interpretation of the local, lek-level results is relatively straightforward. The areal disturbance from OAG is a well supported strongly negative predictor of male attendance at individual leks. The effect is much stronger at a lag of one year and when considering wells within a radius of 3.2 km. However, this should not be taken to imply there are no delayed effects nor no disturbances from more distant wells. There is little uncertainty in the magnitude of the local declines - an areal disturbance of 3% is associated with an average decline of 50% while a 6% decline is associated with a decline of 75%. When scaled up, the lek level results suggest that by 2016 the impact of OAG was equivalent to the loss of 20% of the birds in the Northeast and 30% in the Upper Green River.

Interpretation of the population-level results is more complicated. The mean areal disturbance within 3.2 km of all the leks was a weakly supported predictor of the annual density change with a small effect size. Yet, the predicted population-level impacts were consistent with summation of the local impacts. This apparent paradox it due to three statistical phenomena. The first is that when the statistical power is low an important variable can have low predictive value and therefore receive a low Akaike’s weight. The second is that when density dependence is weak a small effect on the expected density change can have a more substantial impact on the long-term carry capacity. The third is that a standardized effect size can provide a misleading summary of the scale of the possible impact if the variation is highly skewed. The first two phenomena are related in that weak density dependence allows large population fluctuations which obscure the inference of relationships and lower the power. All three factors are at play when dealing with sage-grouse in Wyoming.

It may be possible to reduce the uncertainty by modifying the population dynamic model so that it more closely matched the sage-grouse life history (i.e. most males begin lekking in their third year) and/or through the incorporation of additional variables (Ramey et al., 2011). Aside from this the only other option outside of ethically questionable population-level experiments is to expand the analysis to incorporate data from additional populations across the species range. To enable this, sage-grouse lek count data should be made available to researchers by all states and provinces.

### Climatic Variation

The other key conclusion of this paper is that regional climatic variation is responsible for the inter-decadal population fluctuations experienced by sage-grouse in Wyoming. More specifically, the PDO index is an important predictor of changes in sage-grouse numbers at both the lek and population level. This is perhaps unsurprising as the PDO has previously been used, in combination with the Atlantic Multi-Decadal Oscillation and El Nino Southern Oscillation, to predict drought, drought-related fire frequency, and precipitation trends in the western USA and Rocky Mountains (McCabe et al., 2004; Schoennagel et al., 2007; Kitchen, 2015; Heyerdahl et al., 2008).

Although the current study does not identify the causal pathways through which sea surface temperatures in the North Pacific affects the sage-grouse population dynamics we note that in Wyoming, a positive PDO correlates with cooler, wetter weather, while a negative phase tends to produce warmer, drier conditions (McCabe et al., 2004). We also note that given the relatively poor performance of local precipitation and temperature metrics (Blomberg et al., 2012; Green et al., 2016; Blomberg et al., 2014, 2017; Coates et al., 2016; Gibson et al., 2017; Green et al., 2016), the causal pathways may be complex and involve other organisms such as parasites (Cattadori et al., 2005; Taylor et al., 2013). In fact the complexity of such pathways is one of the reasons that large-scale climate indices such as the PDO often outperform local weather data in predicting population dynamics and ecological process (Stenseth et al., 2002; Hallett et al., 2004). Additional studies to assess the explanatory value of the PDO index across the species range are needed (Doherty et al., 2016). It is noteworthy that the effect of the PDO index appears to be stronger in the most recent part of the time series at least at the population-level. It is also worth noting that the strength of the population-level response appears to vary between working groups (ie the fluctuations in Bates Hole are much bigger than those in the Upper Snake River) and may have a latitudinal or altitudinal component.

The finding that the PDO index is an important driver of sage-grouse abundance in Wyoming has major implications for our understanding and conservation of the species. At the very least it is expected that any long-term population trends, like those of songbirds in western North America (Ballard et al., 2003; McClure et al., 2012), will be better understood in the context of the PDO. At best, it should allow regulators to account for and predict (Stenseth et al., 2003) the effects of climatic variation on sage-grouse population fluctuations, and therefore more effectively balance conservation efforts.

## ACKNOWLEDGMENTS

We thank T.J. Christiansen and the State of Wyoming for providing the lek count data. We also thank L. Brown and R.L. Irvine for comments and edits.

## SUPPLEMENTAL

**Figure S1.**
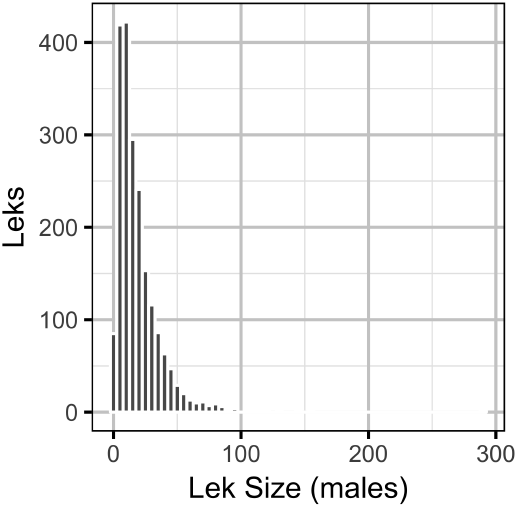
Bayesian estimates of the frequency of leks by count of male sage-grouse in a typical year with no oil and gas.

**Figure S2.**
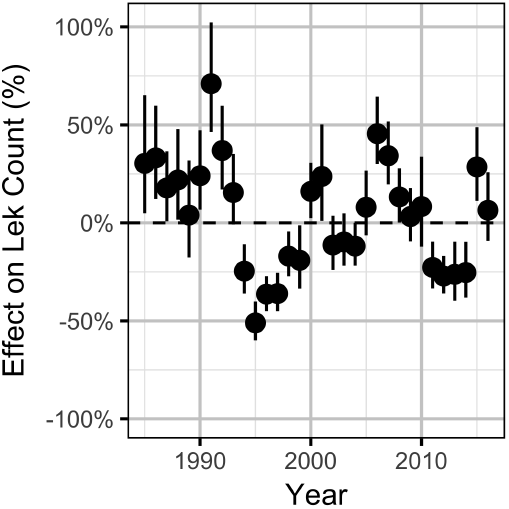
Bayesian estimates (with 95% CIs) of the effect of year on the expected count of male sage-grouse at a typical lek after accounting for the Pacific Decadal Oscillation index and oil and gas.

**Figure S3.**
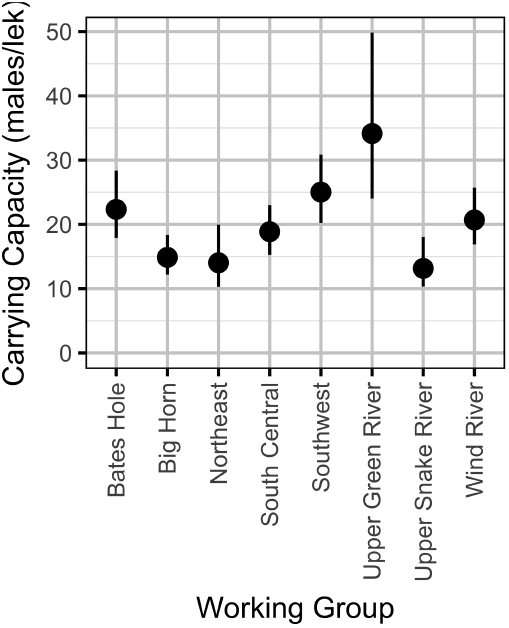
Bayesian estimates (with 95% CIs) of the carrying capacity in a typical year with no oil and gas by working group.

**Figure S4.**
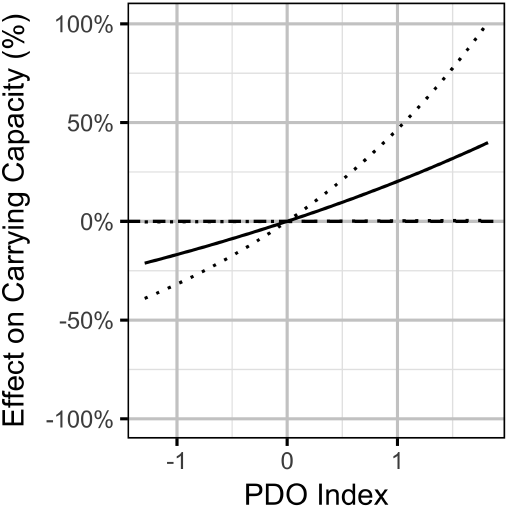
Bayesian estimates (with 95% CIs) of the effect of the Pacific Decadal Oscillation index on the expected carrying capacity at a typical working group. The effect is the percent change in the expected carrying capacity relative to a Pacific Decadal Oscillation index value of 0.

**Figure S5.**
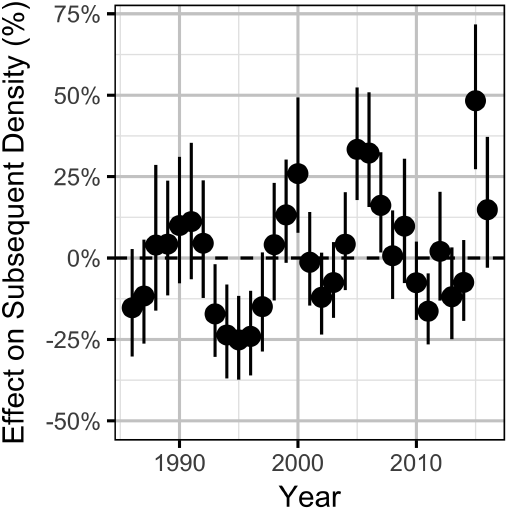
Bayesian estimates (with 95% CIs) of the effect of year on the density the subsequent year after accounting for the Pacific Decadal Oscillation index and oil and gas.

**Figure S6.**
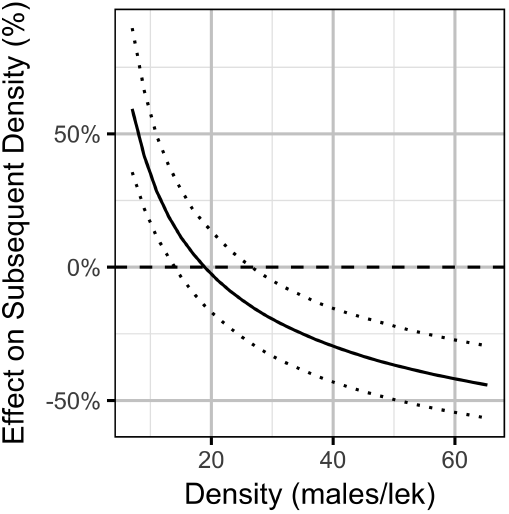
Bayesian estimates (with 95% CIs) of the effect of density on the density the subsequent year with no oil and gas.

**Table S1.** The relative importance (*w_i_*) of spatial scale as a predictor of the count of males sage-grouse at individual leks. The relative importance is across all models with the areal disturbance due to well pads and the Pacific Decadal Oscillation index both independently lagged one to four years.

| Distance (km) | Models | Proportion | $w_i$ |
| --- | --- | --- | --- |
| 3.2 | 16 | 0.25 | 1.00 |
| 1.6 | 16 | 0.25 | 0.00 |
| 6.4 | 16 | 0.25 | 0.00 |
| 0.8 | 16 | 0.25 | 0.00 |

**Table S2.**
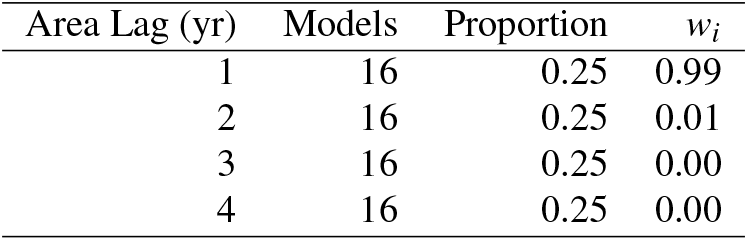
The relative importance (*w_i_*) of the lag in areal disturbance due to well pads as a predictor of the count of males sage-grouse at individual leks. The relative importance is across all models with a lek distance of 0.8, 1.6, 3.2 and 6.4 km and the Pacific Decadal Oscillation index lagged one to four years.

**Table S3.** The model-averaged Maximum Likelihood parameter estimates for the final lek count models with lower and upper 95% CIs. The estimates are for a lek distance of 3.2 km, areal disturbance due to well pads of one year and Pacific Decadal Oscillation index lag of two years.

| Parameter | Estimate | SD | Lower | Upper |
| --- | --- | --- | --- | --- |
| $\beta_A$ | -0.210 | 0.015 | -0.241 | -0.180 |
| $\beta_0$ | 2.491 | 0.058 | 2.378 | 2.605 |
| $\beta_P$ | 0.164 | 0.054 | 0.059 | 0.270 |
| $\log(\phi)$ | -0.908 | 0.014 | -0.935 | -0.882 |
| $\log(\sigma_Y)$ | -1.256 | 0.130 | -1.512 | -1.001 |
| $\log(\sigma_L)$ | -0.011 | 0.019 | -0.048 | 0.026 |

**Table S4.** The Maximum Likelihood parameter estimates for the final full lek count model with lower and upper 95% CIs. The estimates are for a lek distance of 3.2 km, areal disturbance due to well pads of one year and Pacific Decadal Oscillation index lag of two years.

| Parameter | Estimate | SD | Lower | Upper |
| --- | --- | --- | --- | --- |
| $\beta_A$ | -0.210 | 0.015 | -0.241 | -0.180 |
| $\beta_0$ | 2.490 | 0.057 | 2.378 | 2.603 |
| $\beta_P$ | 0.168 | 0.049 | 0.072 | 0.263 |
| $\log(\phi)$ | -0.908 | 0.014 | -0.935 | -0.882 |
| $\log(\sigma_Y)$ | -1.260 | 0.128 | -1.511 | -1.008 |
| $\log(\sigma_L)$ | -0.011 | 0.019 | -0.048 | 0.026 |

**Table S5.** The Bayesian parameter estimates for the final full lek count model with lower and upper 95% CIs. The estimates are for a lek distance of 3.2 km, areal disturbance due to well pads of one year and Pacific Decadal Oscillation index lag of two years.

| Parameter | Estimate | SD | Lower | Upper |
| --- | --- | --- | --- | --- |
| $\beta_A$ | -0.210 | 0.015 | -0.242 | -0.182 |
| $\beta_0$ | 2.488 | 0.062 | 2.370 | 2.615 |
| $\beta_P$ | 0.166 | 0.052 | 0.067 | 0.264 |
| $\log(\phi)$ | -0.908 | 0.013 | -0.934 | -0.881 |
| $\log(\sigma_Y)$ | -1.212 | 0.137 | -1.448 | -0.918 |
| $\log(\sigma_L)$ | -0.006 | 0.019 | -0.044 | 0.030 |

**Table S6.** The model-averaged Maximum Likelihood parameter estimates for the final population models with lower and upper 95% CIs. The estimates are for a lek distance of 3.2 km, areal disturbance due to well pads of one year and Pacific Decadal Oscillation index lag of one year.

| Parameter | Estimate | SD | Lower | Upper |
| --- | --- | --- | --- | --- |
| $\beta_A$ | -0.012 | 0.021 | -0.052 | 0.029 |
| $\beta_D$ | -0.464 | 0.064 | -0.589 | -0.339 |
| $\beta_0$ | 1.351 | 0.179 | 1.000 | 1.702 |
| $\beta_L$ | -0.304 | 0.043 | -0.389 | -0.219 |
| $\beta_P$ | 0.056 | 0.044 | -0.031 | 0.142 |
| $\log(\sigma_Y)$ | -1.692 | 0.156 | -1.997 | -1.386 |
| $\log(\sigma_G)$ | -3.006 | 0.310 | -3.614 | -2.398 |
| $\log(\sigma_\eta)$ | -0.489 | 0.175 | -0.833 | -0.146 |

**Table S7.** The Maximum Likelihood parameter estimates for the final full population model with lower and upper 95% CIs. The estimates are for a lek distance of 3.2 km, areal disturbance due to well pads of one year and Pacific Decadal Oscillation index lag of one year.

| Parameter | Estimate | SD | Lower | Upper |
| --- | --- | --- | --- | --- |
| $\beta_A$ | -0.027 | 0.024 | -0.073 | 0.020 |
| $\beta_D$ | -0.464 | 0.064 | -0.588 | -0.339 |
| $\beta_0$ | 1.352 | 0.178 | 1.002 | 1.702 |
| $\beta_L$ | -0.305 | 0.043 | -0.389 | -0.220 |
| $\beta_P$ | 0.073 | 0.035 | 0.005 | 0.142 |
| $\log(\sigma_Y)$ | -1.708 | 0.152 | -2.005 | -1.410 |
| $\log(\sigma_G)$ | -3.030 | 0.314 | -3.645 | -2.414 |
| $\log(\sigma_\eta)$ | -0.487 | 0.175 | -0.829 | -0.145 |

**Table S8.** The Bayesian parameter estimates for the final full population model with lower and upper 95% CIs. The estimates are for a lek distance of 3.2 km, areal disturbance due to well pads of one year and Pacific Decadal Oscillation index lag of one year.

| Parameter | Estimate | SD | Lower | Upper |
| --- | --- | --- | --- | --- |
| $\beta_A$ | -0.025 | 0.024 | -0.074 | 0.020 |
| $\beta_D$ | -0.468 | 0.065 | -0.594 | -0.345 |
| $\beta_0$ | 1.372 | 0.179 | 1.019 | 1.718 |
| $\beta_L$ | -0.305 | 0.044 | -0.396 | -0.224 |
| $\beta_P$ | 0.073 | 0.037 | 0.001 | 0.144 |
| $\log(\sigma_Y)$ | -1.667 | 0.153 | -1.951 | -1.358 |
| $\log(\sigma_G)$ | -2.907 | 0.353 | -3.592 | -2.154 |
| $\log(\sigma_\eta)$ | -0.483 | 0.180 | -0.788 | -0.094 |

## REFERENCES

Applegate, D. H. and Owens, N. L. (2014). Oil and gas impacts on Wyoming’s sage-grouse: summarizing the past and predicting the foreseeable future. Human-Wildlife Interactions, 8(2):284.

Ballard, G., Geupel, G. R., Nur, N., and Gardali, T. (2003). Long-term declines and decadal patterns in population trends of songbirds in Western North America, 1979–1999. The Condor, 105:737–755.

Blickley, J. L., Blackwood, D., and Patricelli, G. L. (2012a). Experimental Evidence for the Effects of Chronic Anthropogenic Noise on Abundance of Greater Sage-Grouse at Leks: *Greater Sage-Grouse Abundance and Noise*. Conservation Biology, 26(3):461–471.

Blickley, J. L., Word, K. R., Krakauer, A. H., Phillips, J. L., Sells, S. N., Taff, C. C., Wingfield, J. C., and Patricelli, G. L. (2012b). Experimental Chronic Noise Is Related to Elevated Fecal Corticosteroid Metabolites in Lekking Male Greater Sage-Grouse (Centrocercus urophasianus). PLoS ONE, 7(11):e50462.

Blomberg, E. J., Gibson, D., Atamian, M. T., and Sedinger, J. S. (2014). Individual and environmental effects on egg allocations of female Greater Sage-Grouse. The Auk, 131(4):507–523.

Blomberg, E. J., Gibson, D., Atamian, M. T., and Sedinger, J. S. (2017). Variable drivers of primary versus secondary nesting; density-dependence and drought effects on greater sage-grouse. Journal of Avian Biology.

Blomberg, E. J., Sedinger, J. S., Atamian, M. T., and Nonne, D. V. (2012). Characteristics of climate and landscape disturbance influence the dynamics of greater sage-grouse populations. Ecosphere, 3(6):art55.

Blomberg, E. J., Sedinger, J. S., Nonne, D. V., and Atamian, M. T. (2013). Annual male lek attendance influences count-based population indices of greater sage-grouse: Lek Attendance of Greater Sage-Grouse. The Journal of Wildlife Management, 77(8):1583–1592.

Bolker, B. M., Brooks, M. E., Clark, C. J., Geange, S. W., Poulsen, J. R., Stevens, M. H. H., and White, J.-S. S. (2009). Generalized linear mixed models: a practical guide for ecology and evolution. Trends in Ecology & Evolution, 24(3):127–135.

Bradford, M. J., Korman, J., and Higgins, P. S. (2005). Using confidence intervals to estimate the response of salmon populations (Oncorhynchus spp.) to experimental habitat alterations. Canadian Journal of Fisheries and Aquatic Sciences, 62(12):2716–2726.

Braun, C. E., Oedekoven, O. O., and Cameron, L. A. (2002). Oil and Gas Development in Western North America: Effects on Sagebrush Steppe Avifauna with Particular Emphasis on Sage-grouse. In Transactions of the 67th North American Wildlife and Natural Resources Conference, pages 337–349, Washington, D.C. Wildlife Management Institute.

Braun, C. E. and Schroeder, M. A. (2015). Age and sex identification from wings of sage-grouse: Sage-grouse Age and Sex Classification. Wildlife Society Bulletin, 39(1):182–187.

Brooks, S. and Gelman, A. (1998). General methods for monitoring convergence of iterative simulations. Journal of Computational and Graphical Statistics, 7(4):434–455.

Burnham, K. P. and Anderson, D. R. (2002). Model Selection and Multimodel Inference: A Practical Information-Theoretic Approach, volume Second Edition. Springer, New York.

Cattadori, I. M., Haydon, D. T., and Hudson, P. J. (2005). Parasites and climate synchronize red grouse populations. Nature, 433(7027):737–741.

Christiansen, T. J. and Belton, L. R. (2017). Wyoming Sage-Grouse Working Groups: Lessons Learned. Human–Wildlife Interactions, 11(3):274–286.

Claridge-Chang, A. and Assam, P. N. (2016). Estimation statistics should replace significance testing. Nature Methods, 13(2):108–109.

Coates, P. S., Ricca, M. A., Prochazka, B. G., Brooks, M. L., Doherty, K. E., Kroger, T., Blomberg, E. J., Hagen, C. A., and Casazza, M. L. (2016). Wildfire, climate, and invasive grass interactions negatively impact an indicator species by reshaping sagebrush ecosystems. Proceedings of the National Academy of Sciences, 113(45):12745–12750.

Connelly, J. W. and Braun, C. E. (1997). Long-term changes in sage grouse Centrocercus urophasianus populations in western North America. Wildlife Biology, 3:229–234.

Copeland, H. E., Doherty, K. E., Naugle, D. E., Pocewicz, A., and Kiesecker, J. M. (2009). Mapping Oil and Gas Development Potential in the US Intermountain West and Estimating Impacts to Species. PLoS ONE, 4(10):e7400.

Copeland, H. E., Pocewicz, A., Naugle, D. E., Griffiths, T., Keinath, D., Evans, J., and Platt, J. (2013). Measuring the Effectiveness of Conservation: A Novel Framework to Quantify the Benefits of Sage-Grouse Conservation Policy and Easements in Wyoming. PLoS ONE, 8(6):e67261.

Davis, A. J., Hooten, M. B., Phillips, M. L., and Doherty, P. F. (2014). An integrated modeling approach to estimating Gunnison sage-grouse population dynamics: combining index and demographic data. Ecology and Evolution, pages n/a–n/a.

Dennis, B., Ponciano, J. M., Lele, S. R., Taper, M. L., and Staples, D. F. (2006). Estimating density dependence, process noise, and observation error. Ecological Monographs, 76(3):323–341.

Doherty, K. E., Evans, J. S., Coates, P. S., Juliusson, L. M., and Fedy, B. C. (2016). Importance of regional variation in conservation planning: a rangewide example of the Greater Sage-Grouse. Ecosphere, 7(10):e01462.

Doherty, K. E., Naugle, D. E., and Evans, J. S. (2010). A Currency for Offsetting Energy Development Impacts: Horse-Trading Sage-Grouse on the Open Market. PLoS ONE, 5(4):e10339.

Dzialak, M. R., Olson, C. V., Harju, S. M., Webb, S. L., Mudd, J. P., Winstead, J. B., and Hayden-Wing, L. (2011). Identifying and Prioritizing Greater Sage-Grouse Nesting and Brood-Rearing Habitat for Conservation in Human-Modified Landscapes. PLoS ONE, 6(10):e26273.

Fedy, B. C. and Aldridge, C. L. (2011). The importance of within-year repeated counts and the influence of scale on long-term monitoring of sage-grouse. The Journal of Wildlife Management, 75(5):1022–1033.

Fedy, B. C., Aldridge, C. L., Doherty, K. E., O’Donnell, M., Beck, J. L., Bedrosian, B., Holloran, M. J., Johnson, G. D., Kaczor, N. W., Kirol, C. P., Mandich, C. A., Marshall, D., McKee, G., Olson, C., Swanson, C. C., and Walker, B. L. (2012). Interseasonal movements of greater sage-grouse, migratory behavior, and an assessment of the core regions concept in Wyoming. The Journal of Wildlife Management, 76(5):1062–1071.

Fedy, B. C. and Doherty, K. E. (2011). Population cycles are highly correlated over long time series and large spatial scales in two unrelated species: greater sage-grouse and cottontail rabbits. Oecologia, 165(4):915–924.

Fodrie, F. J., Able, K. W., Galvez, F., Heck, K. L., Jensen, O. P., López-Duarte, P. C., Martin, C. W., Turner, R. E., and Whitehead, A. (2014). Integrating organismal and population responses of estuarine fishes in Macondo spill research. BioScience, 64(9):778–788.

Fremgen, A. L., Hansen, C. P., Rumble, M. A., Gamo, R. S., and Millspaugh, J. J. (2016). Male greater sage-grouse detectability on leks: Sightability of Male Sage-Grouse on Leks. The Journal of Wildlife Management, 80(2):266–274.

Fukami, T. and Wardle, D. A. (2005). Long-term ecological dynamics: reciprocal insights from natural and anthropogenic gradients. Proceedings of the Royal Society B: Biological Sciences, 272(1577):2105–2115.

Gamo, R. S. and Beck, J. L. (2017). Effectiveness of Wyoming’s Sage-Grouse Core Areas: Influences on Energy Development and Male Lek Attendance. Environmental Management, 59(2):189–203.

Garton, E. O., Connelly, J. W., Horne, J. S., Hagen, C. A., Moser, A., and Schroeder, M. A. (2011). Greater Sage-Grouse Population Dynamics and Probability of Persistence. In Knick, S. T. and Connelly, J. W., editors, Greater sage-grouse: ecology and conservation of a landscape species and its habitats, number no. 38 in Studies in avian biology, pages 293–374. University of California Press, Berkeley, Calif.

Garton, E. O., Wells, A. G., Baumgardt, J. A., and Connelly, J. W. (2015). Greater Sage-Grouse Population Dynamics and Probability of Persistence. Technical report, Pew Charitable Trusts.

Gelman, A., Carlin, J. B., Stern, H. S., Dunson, D. B., Vehtari, A., and Rubin, D. B. (2014). Bayesian data analysis. Chapman & Hall/CRC texts in statistical science. CRC Press, Boca Raton, third edition edition.

Gibson, D., Blomberg, E. J., Atamian, M. T., and Sedinger, J. S. (2017). Weather, habitat composition, and female behavior interact to modify offspring survival in Greater Sage-Grouse. Ecological Applications, 27(1):168–181.

Gill, J. A., Norris, K., and Sutherland, W. J. (2001). Why behavioural responses may not reflect the population consequences of human disturbance. Biological Conservation, 97(2):265–268.

Green, A. W., Aldridge, C. L., and O’donnell, M. S. (2016). Investigating impacts of oil and gas development on greater sage-grouse: Oil and Gas Impacts on Sage-Grouse. The Journal of Wildlife Management.

Gregory, A. J. and Beck, J. L. (2014). Spatial Heterogeneity in Response of Male Greater Sage-Grouse Lek Attendance to Energy Development. PLoS ONE, 9(6):e97132.

Greven, S. and Kneib, T. (2010). On the behaviour of marginal and conditional AIC in linear mixed models. Biometrika, page asq042.

Hallett, T. B., Coulson, T., Pilkington, J. G., Clutton-Brock, T. H., Pemberton, J. M., and Grenfell, B. T. (2004). Why large-scale climate indices seem to predict ecological processes better than local weather. Nature, 430(6995):71–75.

Harju, S. M., Dzialak, M. R., Taylor, R. C., Hayden-Wing, L. D., and Winstead, J. B. (2010). Thresholds and Time Lags in Effects of Energy Development on Greater Sage-Grouse Populations. Journal of Wildlife Management, 74(3):437–448.

Heyerdahl, E. K., Morgan, P., and Riser, J. P. (2008). Multi-season climate synchronized historical fires in dry forests (1650–1900), northern Rockies, USA. Ecology, 89(3):705–716.

Hilborn, R., Bue, B. G., and Sharr, S. (1999). Estimating spawning escapements from periodic counts: a comparison of methods. Canadian Journal of Fisheries and Aquatic Sciences, 56(5):888–896.

Holloran, M. J. (2005). Greater sage-grouse (Centrocercus urophasianus) population response to natural gas field development in western Wyoming.

Holloran, M. J. and Anderson, S. H. (2005). Spatial Distribution of Greater Sage-Grouse Nests in Relatively Contiguous Sagebrush Habitats. The Condor, 107(4):742.

Holloran, M. J., Kaiser, R. C., and Hubert, W. A. (2010). Yearling Greater Sage-Grouse Response to Energy Development in Wyoming. Journal of Wildlife Management, 74(1):65–72.

Johnson, D. and Rowland, M. (2007). The utility of lek counts for monitoring greater sage-grouse. In Reese, K. and Bowyer, R., editors, Monitoring populations of sage-grouse, pages 15–23. University of Idaho, Moscow, Idaho.

Kitchen, S. G. (2015). Climate and human influences on historical fire regimes (AD 1400–1900) in the eastern Great Basin (USA). The Holocene, page 0959683615609751.

Knape, J. and de Valpine, P. (2012). Are patterns of density dependence in the Global Population Dynamics Database driven by uncertainty about population abundance?: Density dependence in the GPDD. Ecology Letters, 15(1):17–23.

Knick, S. T. and Connelly, J. W., editors (2011). Greater sage-grouse: ecology and conservation of a landscape species and its habitats. Number no. 38 in Studies in avian biology. University of California Press, Berkeley, Calif.

Kristensen, K., Nielsen, A., Berg, C. W., Skaug, H., and Bell, B. M. (2016). TMB: Automatic Differentiation and Laplace Approximation. Journal of Statistical Software, 70(5).

Kvasnes, M. A. J., Storaas, T., Pedersen, H. C., Bjørk, S., and Nilsen, E. B. (2010). Spatial dynamics of Norwegian tetraonid populations. Ecological Research, 25(2):367–374.

Lindström, J., Ranta, E., Lindén, H., Lindstrom, J., and Linden, H. (1996). Large-Scale Synchrony in the Dynamics of Capercaillie, Black Grouse and Hazel Grouse Populations in Finland. Oikos, 76(2):221.

Ludwig, G. X., Alatalo, R. V., Helle, P., Linden, H., Lindstrom, J., and Siitari, H. (2006). Short- and long-term population dynamical consequences of asymmetric climate change in black grouse. Proceedings of the Royal Society B: Biological Sciences, 273(1597):2009–2016.

Lukacs, P. M., Burnham, K. P., and Anderson, D. R. (2010). Model selection bias and Freedman’s paradox. Annals of the Institute of Statistical Mathematics, 62(1):117–125.

Lyon, A. G. and Anderson, S. H. (2003). Potential gas development impacts on sage grouse nest initiation and movement. Wildlife Society Bulletin, 31(2):486–491.

Mantua, N. J., Hare, S. R., Zhang, Y., Wallace, J. M., and Francis, R. C. (1997). A Pacific Interdecadal Climate Oscillation with Impacts on Salmon Production. Bulletin of the American Meteorological Society, 78(6):1069–1079.

Maunder, M. N., Deriso, R. B., and Hanson, C. H. (2015). Use of state-space population dynamics models in hypothesis testing: advantages over simple log-linear regressions for modeling survival, illustrated with application to longfin smelt (Spirinchus thaleichthys). Fisheries Research, 164:102–111.

McCabe, G. J., Palecki, M. A., and Betancourt, J. L. (2004). Pacific and Atlantic Ocean influences on multidecadal drought frequency in the United States. Proceedings of the National Academy of Sciences, 101(12):4136–4141.

McClure, C. J. W., Rolek, B. W., McDonald, K., and Hill, G. E. (2012). Climate change and the decline of a once common bird: Climate Change and Blackbird Decline. Ecology and Evolution, 2(2):370–378.

Millar, R. B. (2011). Maximum likelihood estimation and inference: with examples in R, SAS, and ADMB. Statistics in practice. Wiley, Chichester, West Sussex. OCLC: ocn690090037.

Monroe, A. P., Edmunds, D. R., and Aldridge, C. L. (2016). Effects of lek count protocols on greater sage-grouse population trend estimates: Lek Count Timing and Trend Estimates. The Journal of Wildlife Management, 80(4):667–678.

Moran, P. A. P. (1952). The Statistical Analysis of Game-Bird Records. The Journal of Animal Ecology, 21(1):154.

Moran, P. A. P. (1954). The Statistical Analysis of Game-Bird Records. II. The Journal of Animal Ecology, 23(1):35.

Neilson, R., Lenihan, J., Bachelet, D., and Drapek, R. (2005). Climate change implications for sagebrush ecosystems. In Transactions of the 70th North American Wildlife and Natural Resources Conference, pages 145–149.

Pianosi, F., Beven, K., Freer, J., Hall, J. W., Rougier, J., Stephenson, D. B., and Wagener, T. (2016). Sensitivity analysis of environmental models: A systematic review with practical workflow. Environmental Modelling & Software, 79:214–232.

R Core Team (2017). R: A Language and Environment for Statistical Computing.

Ramey, R., Brown, L., and Blackgoat, F. (2011). Oil and gas development and greater sage grouse (Centrocercus urophasianus): a review of threats and mitigation measures. The Journal of Energy and Development, 35(1):49–78.

Ranta, E., Lindstrom, J., and Linden, H. (1995). Synchrony in Tetraonid Population Dynamics. The Journal of Animal Ecology, 64(6):767.

Ross, B. E., Haukos, D., Hagen, C., and Pitman, J. (2016). The relative contribution of climate to changes in lesser prairie-chicken abundance. Ecosphere, 7(6):e01323.

Schoennagel, T., Veblen, T. T., Kulakowski, D., and Holz, A. (2007). Multidecadal climate variability and climate interactions affect subalpine fire occurrence, western Colorado (USA). Ecology, 88(11):2891–2902.

Selås, V., Sonerud, G. A., Framstad, E., Kålås, J. A., Kobro, S., Pedersen, H. B., Spidsø, T. K., and Wiig, O. (2011). Climate change in Norway: warm summers limit grouse reproduction. Population Ecology, 53(2):361–371.

Stan Development Team (2016). RStan: the R interface to Stan.

Stenseth, N. C., Mysterud, A., Ottersen, G., Hurrell, J. W., Chan, K.-S., and Lima, M. (2002). Ecological effects of climate fluctuations. Science, 297(5585):1292–1296.

Stenseth, N. C., Ottersen, G., Hurrell, J. W., Mysterud, A., Lima, M., Chan, K.-S., Yoccoz, N. G., and Adlandsvik, B. (2003). Studying climate effects on ecology through the use of climate indices: the North Atlantic Oscillation, El Nino Southern Oscillation and beyond. Proceedings of the Royal Society of London. Series B: Biological Sciences, 270(1529):2087–2096.

Taylor, R. L., Tack, J. D., Naugle, D. E., and Mills, L. S. (2013). Combined Effects of Energy Development and Disease on Greater Sage-Grouse. PLoS ONE, 8(8):e71256.

Trenberth, K. E. and Hurrell, J. W. (1994). Decadal atmosphere-ocean variations in the Pacific. Climate Dynamics, 9(6):303–319.

Turek, D. (2015). Comparison of the Frequentist MATA Confidence Interval with Bayesian Model-Averaged Confidence Intervals. Journal of Probability and Statistics, 2015:1–9.

Vaida, F. and Blanchard, S. (2005). Conditional Akaike information for mixed-effects models. Biometrika, 92(2):351–370.

Viterbi, R., Imperio, S., Alpe, D., Bosser-peverelli, V., and Provenzale, A. (2015). Climatic control and population dynamics of black grouse (*Tetrao tetrix*) in the Western Italian Alps: Population Dynamics of Alpine Black Grouse. The Journal of Wildlife Management, 79(1):156–166.

Walker, B. L., Naugle, D. E., and Doherty, K. E. (2007). Greater Sage-Grouse Population Response to Energy Development and Habitat Loss. Journal of Wildlife Management, 71(8):2644–2654.

Walsh, D. P., White, G. C., Remington, T. E., and Bowden, D. C. (2004). Evaluation of the lek-count index for greater sage-grouse. Wildlife Society Bulletin, 32(1):56–68.

Wiens, J. A. (1989). Spatial Scaling in Ecology. Functional Ecology, 3(4):385.

